# Mitochondrial damage triggers therapy-induced senescence

**DOI:** 10.1101/2024.12.20.629667

**Authors:** Chrysiida Baltira, Ferhat Alkan, Isabel Mayayo-Peralta, Miriam Guillen-Navarro, Bram Thijssen, Rose Lempers, Kimberley Pello, Daria M. Fedorushkova, Anna Their, Fleur Jochems, Kicky Rozing, Onno B. Bleijerveld, Hans Janssen, Laura E. Kuil, Jos H. Beijnen, William J. Faller, Joana Silva, Rene Bernards, Olaf van Tellingen, Mark C. de Gooijer

**Affiliations:** Division of Pharmacology, The Netherlands Cancer Institute; Plesmanlaan 121, 1066 CX Amsterdam, The Netherlands; Division of Oncogenomics, The Netherlands Cancer Institute; Plesmanlaan 121, 1066 CX Amsterdam, The Netherlands; Division of Molecular Carcinogenesis, Oncode Institute; The Netherlands Cancer Institute; Plesmanlaan 121, 1066 CX Amsterdam, The Netherlands; Proteomics Facility; The Netherlands Cancer Institute; Plesmanlaan 121, 1066 CX Amsterdam, The Netherlands; Electron Microscopy Facility; The Netherlands Cancer Institute; Plesmanlaan 121, 1066 CX Amsterdam, The Netherlands; Department of Pharmacy & Pharmacology; The Netherlands Cancer Institute; Plesmanlaan 121, 1066 CX Amsterdam, The Netherlands; Department of Pharmaceutical Sciences, Utrecht University; Universiteitsweg 99, Utrecht, the Netherlands; Mouse Cancer Clinic, The Netherlands Cancer Institute; Plesmanlaan 121, 1066 CX Amsterdam, The Netherlands; Faculty of Biology, Medicine and Health, Division of Cancer Sciences, University of Manchester; 46 Grafton Street, Manchester, UK

## Abstract

Glioblastoma (GBM) is a fatal brain tumor with a critical need for better therapies. It is known that the PI3K, MAPK, and CDK4/6 signaling pathways are hyper-activated in these tumors; however, previous studies have used very high concentration of inhibitors to assess their importance, with mixed results. Here we developed PMCi, a combination approach that targets all three pathways simultaneously, at clinically-relevant doses. PMCi effectively suppresses GBM cell proliferation in vitro and in vivo, and outperforms monotherapies and dual combinations. PMCi acts by inducing cellular senescence, which is mediated solely by the mitochondria, and, unlike other forms of senescence, is independent of nuclear damage. This phenotype is caused by a reactive oxygen species (ROS)\cGAS-STING\senescence-associated secretory phenotype (SASP) signaling cascade, that acts in a paracrine manner to establish and maintain senescence. Our results demonstrate that mitochondrial damage is sufficient to drive senescence, and that this can be leveraged to target GBM cells.

## INTRODUCTION

Cellular senescence is a biological process characterized by a durable growth arrest ^1^. Senescent cells undergo distinct morphological and functional changes, and release various immunomodulatory factors, collectively termed the senescence-associated secretory phenotype (SASP). Senescence has mostly been studied in non-transformed cells. In these cells senescence acts as a protective measure against oncogenic transformation by halting the replication of damaged or mutated cells. Conversely however, when these senescent cells are not cleared, the SASP can also contribute to tissue dysfunction, inflammation, and cancer occurrence ^2^. In cancer, numerous compounds with diverse mechanisms of action have been identified to promote therapy-induced senescence (TIS). Due its durable effects on cancer cell proliferation, TIS is now being explored as a therapeutic strategy for many cancers. A common trait of TIS is the role of nuclear DNA damage in senescence signaling ^3^. While many cases involve the use of non-targeted chemotherapeutic agents such as temozolomide ^4^ or doxorubicin ^5^, there are relatively few examples of targeted drugs that effectively induce senescence ^6^.

Mitochondria, traditionally known for their pivotal role in cellular energy production, are increasingly recognized as important regulators maintaining the senescent state ^7^. Senescent cells uphold metabolic activity while harboring dysfunctional mitochondria ^8^. The link between mitochondrial dysfunction and the SASP has recently garnered significant attention. Mitochondrial dysfunction can instigate the release of mitochondrial DNA (mtDNA) into the cytoplasm, thereby eliciting an inflammatory response and triggering SASP activation ^9,10^. This process is mediated by the cGAS-STING pathway, which recognizes cytoplasmic DNA and initiates a signaling cascade leading to the production of pro-inflammatory cytokines. Notably, in the context of nuclear DNA damage the influence of mitochondrial dysfunction and mtDNA release seems to be specifically related to the SASP feature of the senescent state as it does not affect the senescence hallmark feature of proliferation arrest.

While existing studies provide valuable insights, there remains a gap in understanding the role of mitochondria in TIS of cancerous cells, especially in the context of targeted anticancer treatment. Here we utilize a triple-target combination therapy, combining inhibitors targeting the PI3K, MAPK and CDK4/6 pathways (PMCi). These pathways are concurrently over-activated in the vast majority of glioblastoma (GBM) patients (Fig. 1A) ^11^, making simultaneous inhibition of all three pathways a logical therapeutic avenue for GBM albeit one that has so far remained unexplored due the complexity of polypharmacology. Strikingly, we find that PMCi not just efficiently inhibits cell proliferation, but also induces TIS. PMCi treatment in GBM induces senescence independently of nuclear DNA damage, but is notably driven by dysregulated mitochondria. We find that these dysfunctional mitochondria not only contribute to the production of the SASP, but they also play a central role in triggering senescence through mitochondrial oxidative damage. Together, these results describe a previously unappreciated phenomenon in which dysfunctional mitochondria are required and sufficient to drive therapy-induced senescence (mtTIS).

**Fig. 1.**
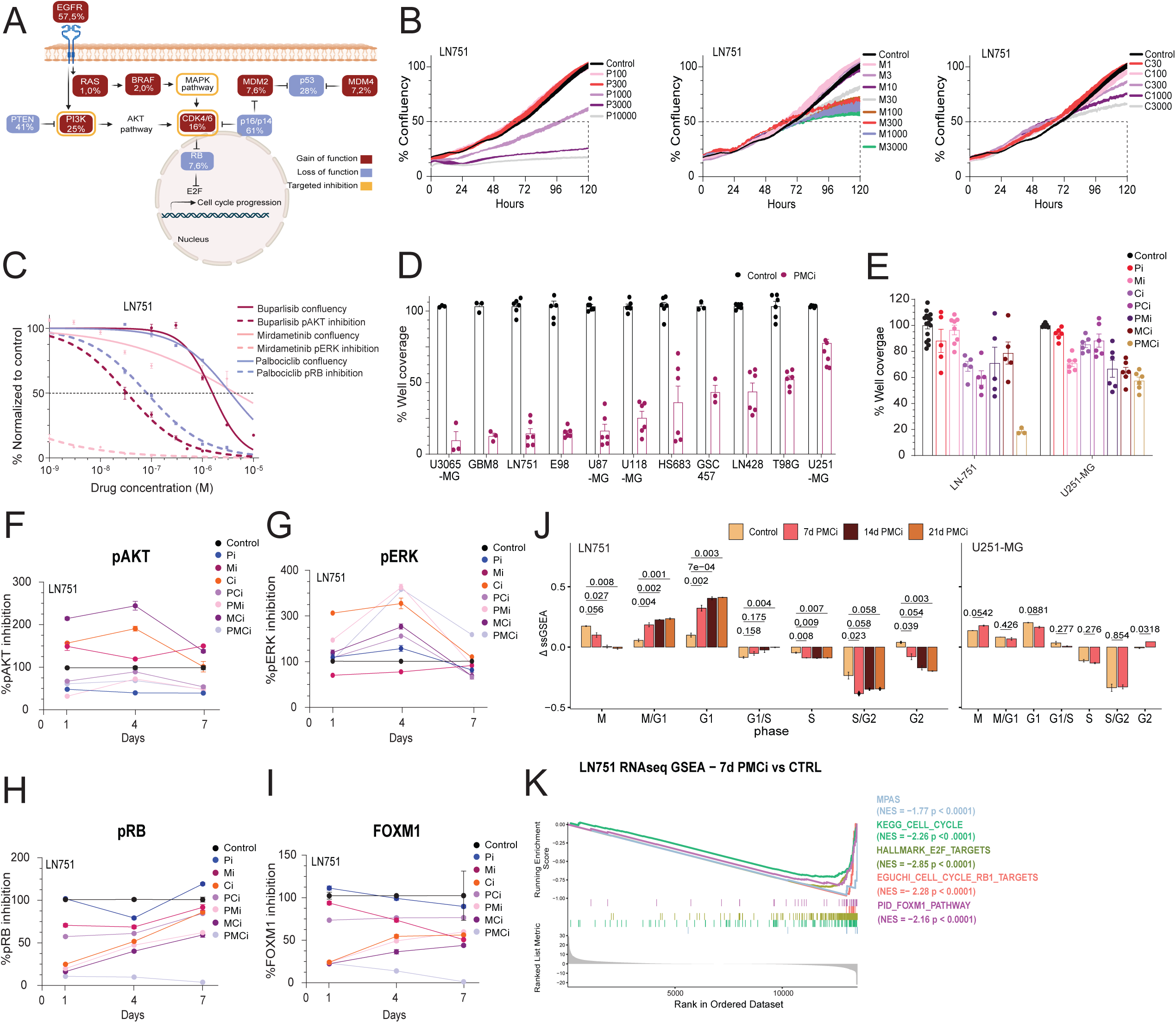
Low doses of PMCi efficiently inhibit proliferation of GBM cells by durable inhibition of the CDK4/6-Rb pathway. **(A)** Diagram illustrating common pathway alterations in GBM patients and the corresponding targets of PMCi treatment. created in BioRender.com. **(B)** Incucyte data for the LN751 cell line showing percentage of confluency normalized to control samples (at 120 h) across various concentrations of P, M, and C inhibitors. n = 2 independent experiments, with line thickness representing SEM **(C)** Solid lines show drug concentrations (P, M or C) versus proliferation (confluency data from panel B at 120 h). Dashed lines show corresponding p-AKT, p-ERK and p-RB levels assessed by Simple Western **(D)** Proliferation (%well coverage normalized to controls) across a panel of cell lines treated for 7 days with PMCi concentrations fixed at 300 nM Buparlisib, 100 nM Mirdametinib, and 30 nM Palbociclib. Minimum n = 2 independent experiments **(E)** Proliferation of LN751 and U251-MG treated with single, double, and triple combinations of PMCi. Minimum n = 2 independent experiments. **(F)** Simple Western analyses showing p-AKT, **(G)** p-ERK, **(H)** p-RB, and **(I)** FOXM1 levels in LN751 cells treated with single, double, and triple PMCi combinations at 1, 4, and 7 days. **(J)** Cell cycle stage scores for PMCi-treated and untreated cells, derived from the single-sample Gene Set Enrichment Analysis (ssGSEA) of RNAseq data, showing the sample-specific ΔssGSEA scores (difference from mean) for 7 genesets associated with different cell-cycle stages. **(K)** GSEA comparing cells treated with PMCi for 7 days to control cells in the responsive LN751 cell lines. The plot shows normalized enrichment scores (NES) for various gene sets, with different colors indicating distinct gene sets and the corresponding p values. Error bars represent mean ± SEM.

## RESULTS

### PMCi efficiently inhibits GBM cell proliferation by inducing senescence

We investigated the potency of targeting the PI3K, MAPK, and CDK4/6 pathways in GBM cell lines using the inhibitors buparlisib, mirdametinib, and palbociclib. We first assessed the impact of monotherapy on cell proliferation across three different GBM cell lines, using the Incucyte system. Cell confluency was only more than 50% reduced at concentrations exceeding 1000 nM in all three cell lines (Fig. 1B, Fig. S1A). We further explored target inhibition by examining their downstream effectors p-Akt, p-ERK, and p-RB. Notably, Simple Western analysis after a 3-hour treatment revealed that target inhibition occurred at concentrations manifold lower than the IC obtained by the Incucyte system (Fig. 1C and Fig. S1A). This effect was most pronounced for inhibition of p-ERK (0.2 nM *vs*. 4400 nM). These findings highlight a disconnect between target inhibition and the resulting proliferation response. We hypothesize that single-dose drugs are effective only at high concentrations, where they exert off-target effects that influence multiple pathways involved in proliferation. This is consistent with the known concurrent over-activation of various cell proliferation pathways in GBM, which facilitates rapid pathway rewiring and sustained proliferation.

Consequently, we sought to investigate a triple drug combination, administered at considerably lower concentrations, to assess whether it could more effectively inhibit proliferation compared to single-agent treatments. We initiated our investigation using a combination of 300 nM buparlisib (PI3K inhibitor-Pi), 100 nM mirdametinib (MEK inhibitor - Mi), and 30 nM palbociclib (CDK4/6 inhibitor - Ci) across a panel of GBM cell lines (Fig. 1D). The sensitivity to this combination (from now on referred to as PMCi) varied across cell lines, with some being highly sensitive, while others, such as U251-MG, showing only minimal proliferation inhibition. Next, we screened a broad range of concentrations of the three drugs as single agent, double and triple combinations in two serum-cultured human GBM cell lines and two human glioma stem cell (GSC) lines. PMCi consistently demonstrated superior efficacy over monotherapies and dual-drug combinations in suppressing proliferation (Fig. 1E and fig. S1B-C). Notably, U251-MG cells only showed inhibition at very high MCi concentrations, far exceeding those required for single-agent target inhibition (fig. S1D), confirming its relative resistance. As a result, we used U251-MG as a model for a non-responsive line in subsequent experiments. To further validate the superiority of PMCi, we confirmed its efficacy in the responsive LN-751 cell line using alternative inhibitors targeting the same pathways (fig. S1E). This demonstrated that PMCi was superior to dual and monotherapy regardless of the inhibitors used.

Focusing on the responsive LN-751 cell line, we evaluated target inhibition of the selected triple PMCi combination (fig. S1C) using Simple Western analysis. Our objective was to understand why the PMCi combination demonstrates superior efficacy compared to dual combinations and monotherapies. As expected, treatment with Pi alone resulted in a significant reduction in the downstream PI3K target p-AKT (Fig. 1F). However, treatment with Mi or Ci alone resulted in markedly elevated p-AKT levels, suggesting compensatory pathway rewiring. Notably, this upregulation of p-AKT was effectively suppressed in all combinations that included Pi (Fig. 1F). As shown before, p-ERK was down by more than 90% following 3-hour exposure to 3 nM of mirdametinib alone (Fig. 1C). However, p-ERK levels began to rebound after 1 day (Fig. 1G), which we determined was not due to drug instability, as confirmed by LC-MS/MS measurements (fig. S1F). Furthermore, p-ERK significantly increased when exposed to Ci, PCi and even PMCi, especially at 4 days (Fig. 1G). Similarly, while Ci initially decreased pRB levels by 80%, a gradual rebound occurred during subsequent days (Fig. 1H). Importantly, this rebound was not prevented by PCi or MCi. Triple PMCi not only prevented this rebound, but actually further declined p-RB to undetectable levels, which is in line with robust proliferation inhibition (Fig. 1H). Moreover, PMCi uniquely induced complete downregulation of FOXM1 (Fig. 1I), a transcription factor regulated by pRB and expressed in cycling cells ^12^, further supporting PMCi efficacy in suppressing proliferation.

In line with the proliferation and target inhibition data, gene set enrichment analysis (GSEA) of RNA-Seq data from the responsive LN-751 line after 7, 14, and 21 days of PMCi treatment revealed a significantly increased G1 population at the expense of all other cell cycle populations (Fig. 1J). This cell cycle inhibition effect was not observed in the less-responsive U251-MG line. Principal component analysis (PCA) confirmed the reproducibility of the samples and showed that the duration of treatment did not affect the response, as these samples clustered together (fig. S1G). GSEA further validated the target inhibition of PMCi, showing significantly reduced enrichment scores for the MAPK Pathway Activity Score, RB targets, E2F targets, FOXM1 targets and the KEGG cell cycle gene set only in the responsive line (Fig. 1K and fig. S1H).

Strikingly, after four days of PMCi we observed that responsive GBM cells started to enlarge and flatten (data not shown). This morphology was full-blown after seven days of PMCi and is indicative of cellular senescence (Fig. 2A and fig. S2A). To validate the induction of senescence across a panel of GBM lines, we employed various techniques and markers. Senescence-associated β-galactosidase (SA-β-gal) positivity was evident (Fig. 2B-C and fig. S2B-C), along with a reduction in the colony potential in drug-free medium following 7 days of exposure to PMCi (Fig. 2D-E and fig. S2D). Additionally, GSEA of the Fridman gene sets (Fig. 2F) and the SENCAN classifier (Fig. 2G), a specific classifier for detecting cancer cell senescence ^13^, confirmed an increase in senescence-associated transcripts. Importantly, none of these features was observed in the non-responsive line U251-MG.

**Fig. 2.**
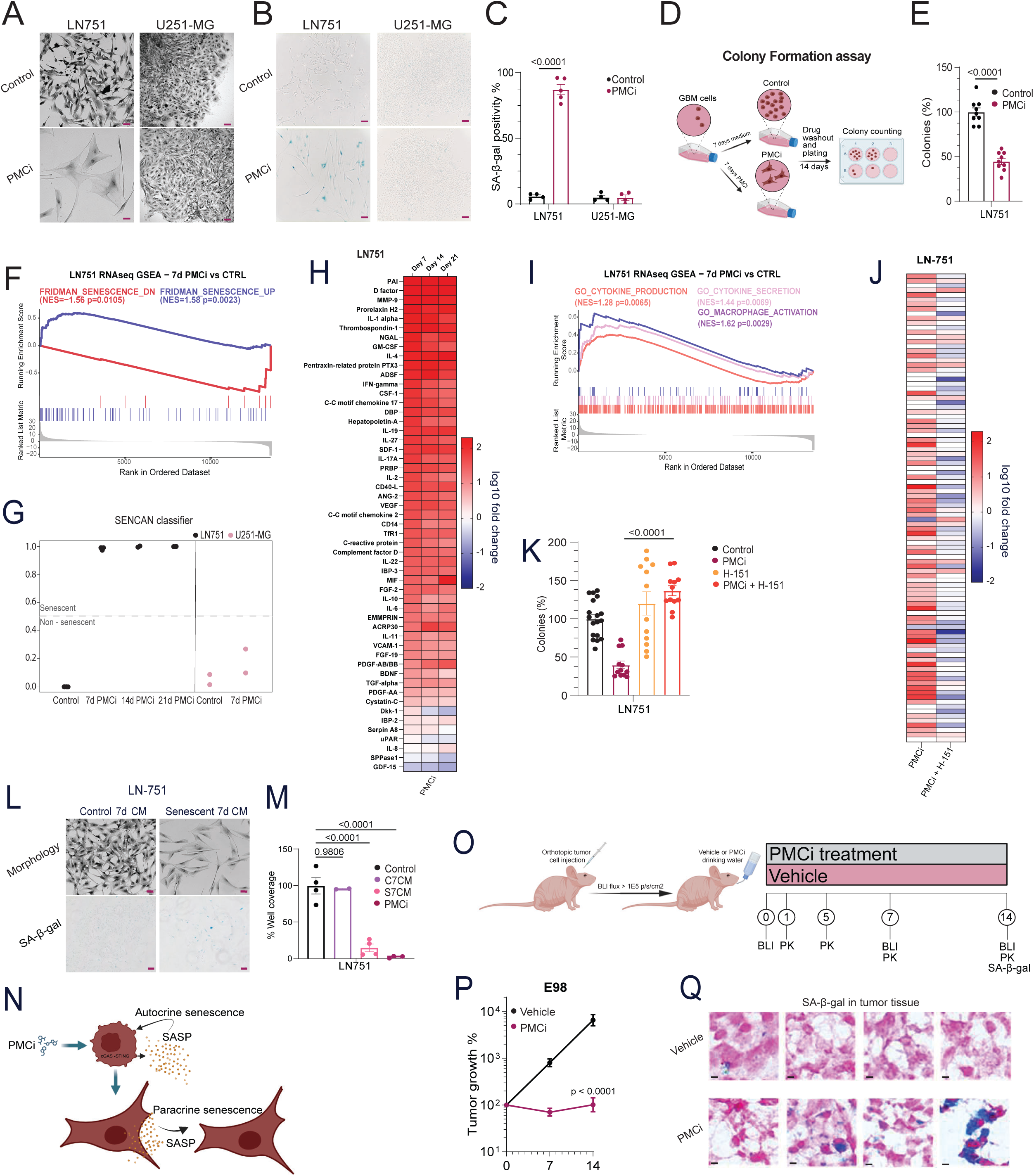
PMCi induces cellular senescence. **(A)** Representative images of morphological changes in GBM cells after 7 days of PMCi treatment (scale bar: 50 µM). **(B)** Representative β-galactosidase (β-gal) staining images of treated cells (scale bar: 100 µM). **(C)** Quantification of β-gal staining in GBM cell lines across two independent experiments (n = 2). Unpaired student t-test (p< 0.005) **(D)** Schematic of the colony formation assay setup, created using BioRender.com. **(E)** Quantification of colony formation following PMCi treatment, based on at least two independent experiments (n ≥ 2).). Unpaired student t-test (p< 0.005) **(F)** Gene Set Enrichment Analysis (GSEA) of RNA-seq data comparing LN-751 cells treated with PMCi for 7 days to control cells, showing normalized enrichment scores (NES) for gene sets with color-coded p-values. **(G)** SENCAN gene expression classifier scores for senescence-sensitive LN-751 and non-responsive U251-MG cell lines, untreated and after PMCi treatment. Scores above the 0.5 threshold (dashed line) indicate senescence. **(H)** Cytokine array heatmaps showing fold changes in SASP factors and immunomodulatory proteins at 7, 14, and 21 days post-PMCi treatment. **(I)** GSEA plot of RNA-seq data in LN-751 cells treated with PMCi for 7 days, displaying NES and p-values for various gene sets. **(J)** Heatmap of cytokine fold changes comparing PMCi-treated cells to cells treated with PMCi plus H151. **(K)** Quantification of colony formation after treatment with PMCi or H151 in two independent experiments. Statistical analysis was conducted using two-way ANOVA with Šidák’s test (P < 0.05). **(L)** Representative images of cell morphology and β-gal staining in cells treated with control conditioned medium (control c.m.) or PMC-conditioned medium (PMC c.m.) for 7 days (scale bars: 50 µM and 100 µM, respectively). **(M)** Quantification of cell coverage in E98 and LN-751 cells treated with various conditioned media or PMCi, analyzed using two-way ANOVA with Šidák’s test (P < 0.05). **(N)** Schematic summarizing findings related to SASP production, created using BioRender.com. **(O)** Schematic of the in vivo experimental procedure. **(P)** Inhibition of E98 intracranial tumor growth by PMCi in Abcg2-/-;Abcb1a/b-/- nude mice, assessed using two-way ANOVA on log-transformed bioluminescence imaging (BLI) data. **(Q)** Representative histochemical β-gal staining of treated and untreated mice. Data are presented as mean ± SEM.

To expand the characterization of senescence-associated features, we next asked whether PMCi- induced senescent GBM cells produced a SASP. Following PMCi treatment and a 24-hour drug washout, conditioned medium was collected for cytokine array analysis, revealing an increased production of key senescence markers IL-1α, IL-6 and IL-8 ^14^ (Fig. 2H and fig. S2E-F). GSEA further revealed a significant enrichment of gene sets comprising cytokine production and secretion, as well as macrophage activation (Fig. 2I). These gene sets, however, were not significantly enriched in the non-responsive U251-MG cell line upon PMCi treatment (fig. S2G). To investigate the underlying regulation of SASP production in PMCi-induced senescent GBM cells, we used the STING inhibitor H151 to examine the involvement of the cGAS-STING pathway, a known inducer of the SASP in fibroblasts ^15–17^. In line with this, co-treatment with PMCi and H151 effectively suppressed SASP production in LN-751 cells compared to cells only exposed to PMCi (Fig. 2J). Strikingly however, it also prevented the PMCi-induced senescence-associated loss of colony potential (Fig. 2K and fig. S2H). The latter effect was more pronounced in LN-751 cells, as H151 alone also reduced the colony formation in E98 cells. To investigate whether the PMCi-induced SASP was functional, we exposed treatment-naïve cells to conditioned medium from PMCi-induced senescent LN-751 and E98 cells. Indeed, we observed that the SASP was capable of inducing paracrine senescence in both cell lines (Fig. 2L-M and fig. S2I-J). These findings together demonstrate that PMCi-treated GBM cells secrete a functional SASP regulated by the cGAS-STING pathway and that the SASP plays a causative role in PMCi-induced senescence (Fig. 2N).

In order to explore PMCi *in vivo*, we orthotopically injected human E98 cells in male and female *Abcg2*^-/-^;*Abcb1a/b*^-/-^ mice and treated them with PMCi for 14 days (Fig. 2O). While buparlisib ^18^ and mirdametinib ^19^ readily penetrate the brain in wild-type mice, this is not the case for palbociclib ^20,21^. However, in mice lacking the dominant efflux transporters at the blood-brain barrier, Abcb1 and Abcg2, the brain penetration of palbociclib is excellent ^20,21^, allowing us to study *in vivo* effects of PMCi in the most pharmacologically favorable setting. We found that PMCi efficiently inhibited tumor growth (Fig. 2P and fig. S3A) and extended survival of mice and fig. S3A-B). Antitumor efficacy was similar between genders (fig. S3C-D). Importantly, we optimized the drug dosages to achieve clinically relevant plasma levels (fig. S3E). These plasma levels did not induce any hematological toxicity after 14 days of exposure (fig. S3F). After harvesting the brains of some of the mice on the last day of treatment, we utilized SA-β-gal staining to assess senescence and found that PMCi could induce SA-β-gal staining in tumor cells (Fig. 2Q), but not in healthy brain tissue and skin (fig. S3G).

### PMCi-induced senescent cells lack nuclear DNA damage but display mitochondrial network and functionality changes

After confirming that PMCi induces senescence, we wanted to understand the underlying mechanism of induction. Nuclear DNA damage is a well-known trigger for cellular senescence and is classically considered a senescence hallmark (26), so we first assessed whether nuclear DNA damage was involved in PMCi-induced senescence. Strikingly, neutral (Fig. 3A) and alkaline comet assays (fig. S4A-B) did not reveal any evidence of single- or double-stranded DNA breaks in PMCi-induced senescent cells. The absence of double-stranded DNA breaks was further corroborated by γH2AX foci analysis (Fig. 3B and fig. S4C). Our results deviate from previous literature reports on cellular senescence induced by CDK4/6 inhibitors such as palbociclib, which typically report DNA damage induction in senescent cells ^22,23^. However, these *in vitro* studies all used palbociclib concentrations of 1000 nM or higher for senescence induction, far exceeding what can be achieved in patients. When using PMCi treatment, senescence occurs at lower, clinically relevant palbociclib concentrations (fig. S1C), Notably, the clinically achievable steady-state plasma concentration is approximately 200 ng/ml, or around 400 nM ^24^. We confirmed that nuclear DNA damage – roughly at level with that incurred by 0.5 Gy of ionizing radiation - occurs at palbociclib concentrations of 1000 nM (Fig. 3B and fig.S4C), yet this finding is most likely an *in vitro* artifact with minimal, if any, clinical relevance.

**Fig. 3.**
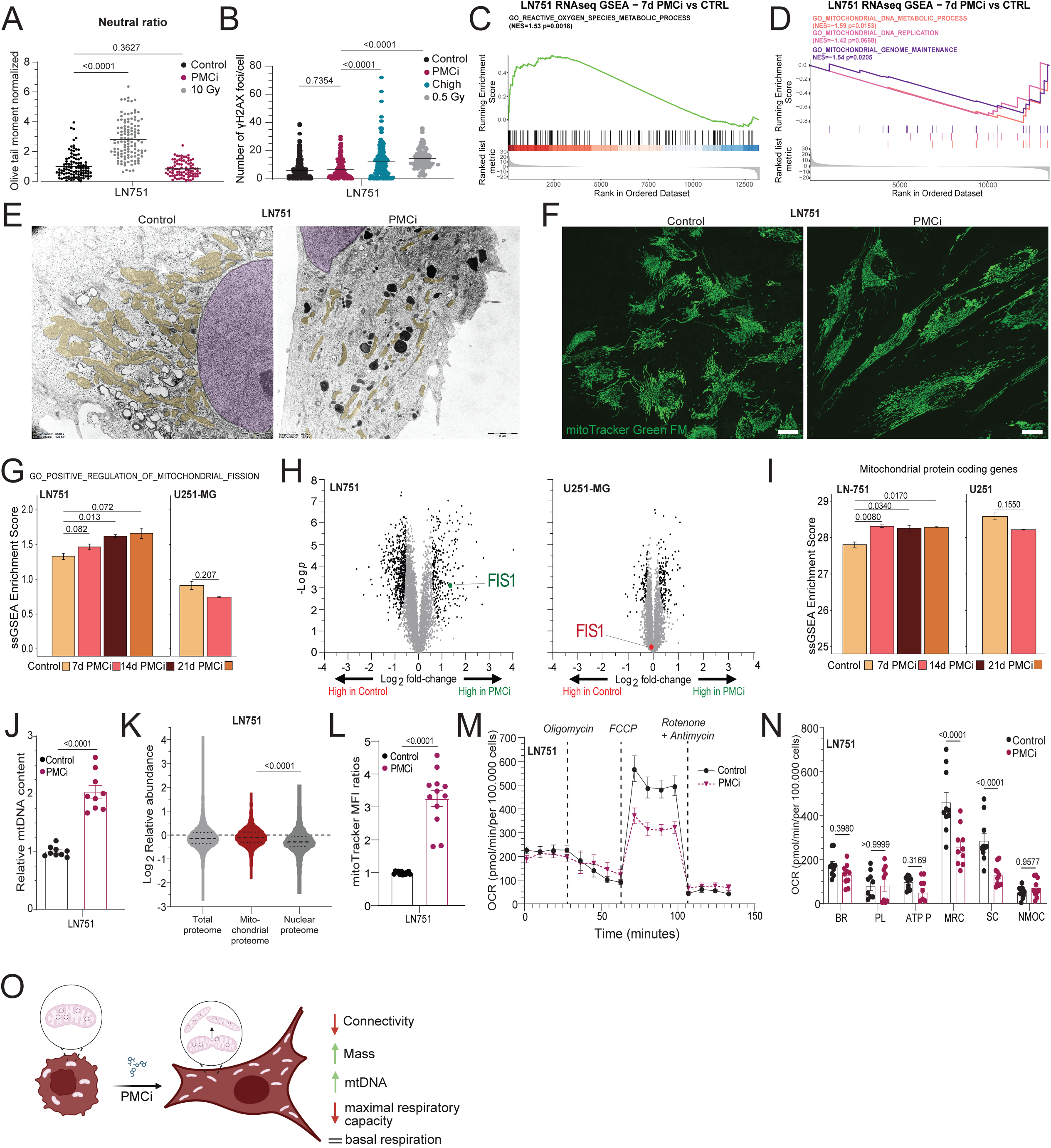
PMCi-induced senescent GBM cells exhibit mitochondrial dysfunctionality in absence of nuclear DNA damage. **(A)** Quantification of neutral comet assay Olive tail moment normalized to control samples in untreated, irradiated, and PMCi-treated cells (n = 2 independent experiments). Data analyzed using one-way ANOVA with Tukey’s multiple comparisons test. **(B)** Quantification of γH2AX foci in LN-751 cells treated with PMCi or 1000 nM Palbociclib (Chigh). Data analyzed with one-way ANOVA and Tukey’s multiple comparisons test. **(C)** Gene Set Enrichment Analysis (GSEA) of RNA-seq data comparing LN-751 cells treated with PMCi for 7 days to untreated controls. **(D)** NES for various gene sets from GSEA, with distinct colors indicating gene sets and corresponding p-values. **(E)** Representative transmission electron microscopy (TEM) images of control and senescent cells highlighting mitochondria (yellow) and nucleus (pink), scale bar = 2 μm. **(F)** Confocal microscopy images of proliferative and senescent cells stained with MitoTracker Green FM to visualize mitochondria (scale bar = 50 μm). **(G)** ΔssGSEA enrichment scores derived from RNA-seq data comparing mitochondrial fission gene sets in PMCi-treated versus untreated cells (n = 1 independent experiment, performed in triplicate). **(H)** Volcano plot from proteomics showing FIS1 changes after 7 days of PMCi treatment, with significant proteins indicated as black dots. **(I)** ΔssGSEA for mitochondrial protein-coding genes (n = 1 independent experiment, performed in triplicate). **(J)** Relative mtDNA-to-nDNA copy number after PMCi treatment quantified via qRT-PCR (n = 3 independent experiments). **(K)** Fold changes in total mitochondrial versus non-mitochondrial proteins post-PMCi treatment, determined by proteomics. **(L)** Mitochondrial mass analyzed using MitoTracker fluorescence and mean fluorescence intensity (MFI) ratios in GBM cells treated with PMCi (n = 3 independent experiments). **(M)** Seahorse X-24 analysis comparing oxygen consumption rate (OCR) in control and PMCi-treated senescent cells, with dotted lines indicating key respiration parameters. **(N)** Quantification of basal respiration (BR), proton leak (PL), ATP production (ATP P), maximal respiratory capacity (MRC), spare capacity (SC), and non-mitochondrial oxygen consumption (NMOC), (n = 3 independent experiments). Data analyzed using two-way ANOVA with Šídák’s multiple comparisons test. **(O)** Schematic of mitochondrial changes in PMCi-induced senescent cells, created using BioRender.com. Statistical analyses for **(G), (I), (J), (K),** and **(L)** were performed using unpaired Student’s t-test. Error bars represent mean ± SEM.

Given the absence of nuclear DNA damage, we explored alternative pathways that might drive SASP production and PMCi-induced senescence. While initial RNA-seq analysis had already highlighted notable alterations in cell cycle-related genes (Fig. 1J-K), we examined our GSEA results further to uncover additional mechanistic clues. The significant deregulation of genes in mitochondria-related GO terms in PMCi-induced senescent LN-751 cells captured our attention, due to mitochondria harboring another major source of cellular DNA. We found an upregulation of genes associated with reactive oxygen species (ROS) production (Fig. 3C) and a downregulation of pathways involved in mitochondrial metabolic processes, mtDNA replication, and mitochondrial genome regulation (Fig. 3D). Notably, these mitochondrial changes were absent in the non-responsive U251-MG line (fig. S4D-E), highlighting the mitochondria as a potential driver of PMCi-induced senescence.

To validate the GSEA findings and characterize mitochondrial function in PMCi-induced senescent cells in greater detail, we first examined the mitochondrial network. Electron microscopy revealed profound alterations in mitochondrial morphology, with mitochondria appearing fragmented and exhibiting an elongated structure (Fig. 3E). These mitochondria also harbored disrupted cristae, characterized by notable gaps, and were more dispersed throughout the cytoplasm, suggesting enhanced fission activity and altered spatial organization (Fig. 3E). Moreover, the distance between the mitochondria and the nucleus increased, further indicating a reorganization of the mitochondrial network. Fluorescence microscopy confirmed these observations, demonstrating a compromised mitochondrial network with reduced interconnectivity (Fig. 3F). Transcriptomic GSEA supported these findings by showing significant enrichment of genes associated with mitochondrial fission (Fig. 3G). Proteomic analysis revealed elevated levels of FIS1, a primary regulator of mitochondrial fission ^25^, corroborating the enhanced fission observed morphologically (Fig. 3H and fig. S4F). Importantly, these disruptions were specific to the PMCi-responsive cell lines LN-751 and E98, with no changes detected in the non-responsive U251-MG cells. Interestingly, while the mitochondrial network was disrupted, PMCi-induced senescent cells harbored significantly increased amounts of mitochondrial gene transcripts (Fig. 3I) and mtDNA copy numbers (Fig. 3J and fig. S4G). Proteomic data indicated a corresponding elevation in the mitochondrial proteome, as evidenced by a comparison of nuclear and mitochondrial proteins (Fig. 3K), consistent with previous work showing superscaling of the mitochondrial proteome in senescent cells ^26^. This was accompanied by an increase in mitochondrial mass (Fig. 3L and fig. S4H), suggesting a potential compensatory upregulation of oxidative phosphorylation (OXPHOS) components. However, despite these compensatory mechanisms, maximal respiratory capacity was significantly reduced in PMCi-induced senescent cells (Fig. 3M-N and fig. S4I-J). Surprisingly, basal respiration remained unchanged in PMCi-induced senescent cells compared to controls. These findings indicate that PMCi-induced senescent cells generate additional mitochondria maintaining basal respiration, yet these newly formed mitochondria are likely less fit and incapable of achieving high respiratory capacity (Fig. 3O).

### PMCi-induced senescence is dependent on mitochondria

The mTOR pathway plays a crucial role in mitochondrial biogenesis ^27^. Given our evidence that PMCi-induced senescent cells are characterized by an increased mitochondrial mass, we hypothesized that mTOR inhibitors could potentially rescue the senescent phenotype. Substantiating our hypothesis, we observed incomplete deactivation of p-AKT despite PI3K inhibition (Fig. 1F). Notably, treatment with a dual mTORC1/2 inhibitor successfully rescued both the altered morphology (Fig. 4A) and the impaired proliferation potential (Fig. 4B and fig. S5A) associated with PMCi-induced senescence. In line with its role in mitochondrial biogenesis, mTOR inhibition effectively attenuated the elevated mitochondrial mass seen in senescent cells (Fig. 4C and fig. S5B). This aligns with prior studies showing that imbalance between cell proliferation and cell growth may lead to cell cycle exit ^26,28–31^. Consistent with these observations, although cell cycle progression is halted by PMCi (Fig. 1), mitochondrial biosynthesis continues (Fig. 3). We therefore hypothesized that this organelle imbalance drives the PMCi-senescent state.

**Fig. 4.**
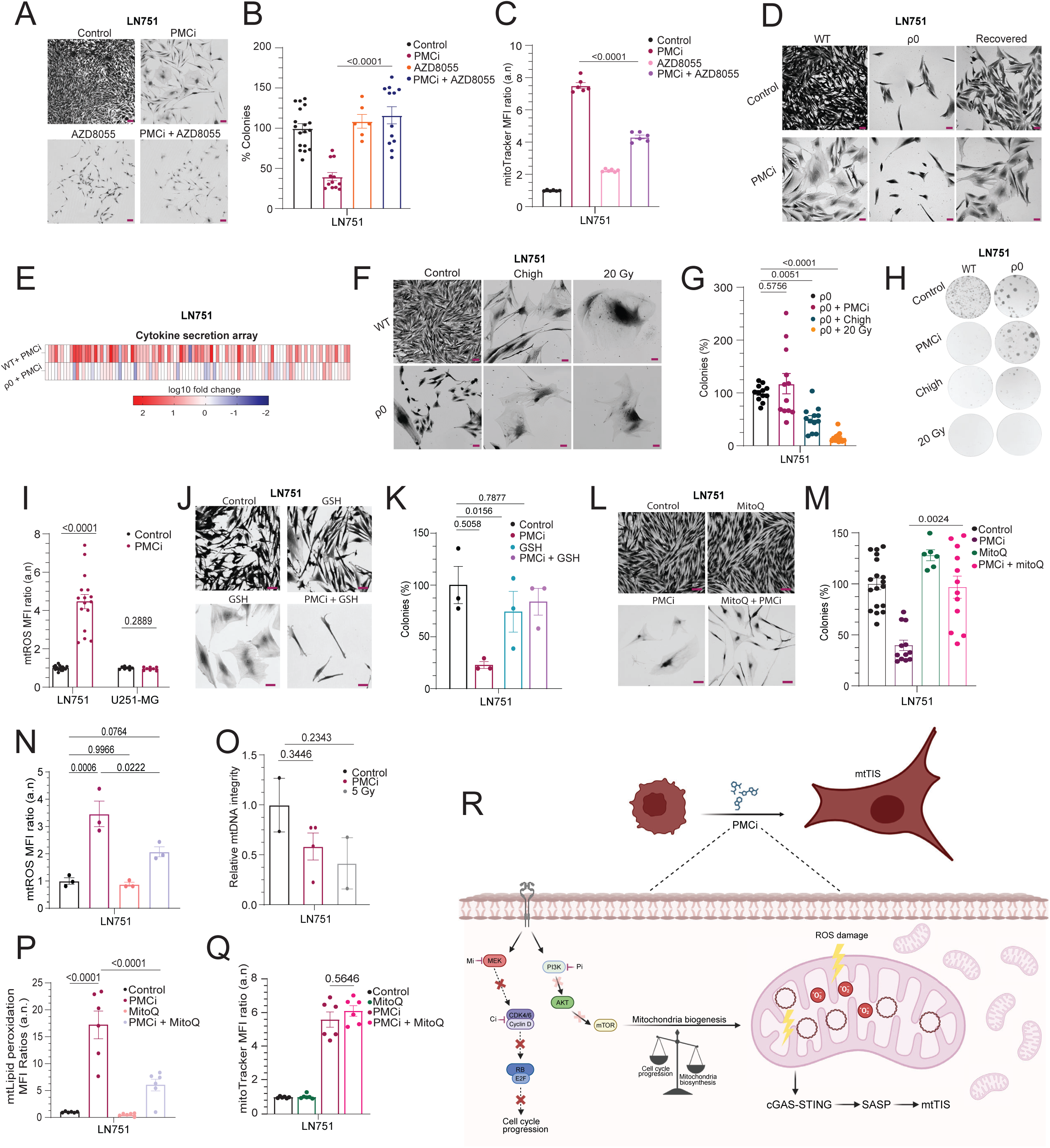
Mitochondrial biogenesis drives PMCi-induced senescence by triggering oxidative mitochondrial damage. **(A)** Representative images of LN751 cell morphology after treatment with PMCi and 100 nM mTOR inhibitor AZD8055 (scale bar: 50 μm; n = 2 independent experiments). **(B)** Colony formation assay under the same treatment conditions, analyzed via flow cytometry (n = 2 independent experiments). **(C)** MitoTracker-based flow cytometry analysis of mitochondrial mass in LN751 cells treated with PMCi and AZD8055, showing median fluorescence intensity (MFI) ratios**. (D)** Representative images of control, ρ0, and recovered ρ0 cells following PMCi treatment. **(E)** Cytokine array heatmap comparing fold changes in cytokine levels between PMCi-treated and untreated WT cells, and PMCi-treated versus untreated ρ0 cells. **(F)** Morphological images of control and ρ0 cells treated with 1000 nM Palbociclib (Chigh) or 20 Gy irradiation (scale bar: 50 μm). **(G)** Colony formation assay for the same conditions, and **(H)** corresponding well plate images stained with crystal violet, comparing WT and ρ0 cells (n = 2 independent experiments). **(I)** MitoSOX-based flow cytometry showing mitochondrial ROS (mtROS) levels as MFI ratios (minimum n = 2 independent experiments). Statistical analysis was performed using an unpaired Student’s t-test. **(J)** Representative images of LN751 cells treated with PMCi, with or without 2 mM glutathione (GSH) antioxidant (scale bar: 50 μm). **(K)** Colony formation assay for LN751 cells treated with PMCi and GSH (n = 1, performed in triplicate). **(L)** Representative images of LN751 cells treated with PMCi, with or without 100 nM mitoQuinone (mitoQ), a mitochondrial antioxidant (scale bar: 50 μm; n = 2 independent experiments). **(M)** Colony formation assay under the same conditions as **(L)** (n = 2 independent experiments). **(N)** MitoSOX-based flow cytometry analysis of mtROS under the same conditions as **(L)**, using MFI ratios (n = 1 experiment, performed in triplicate). **(O)** Relative mtDNA integrity in LN751 cells following PMCi treatment, measured via quantitative PCR (n = 1, with duplicates per condition). **(P)** MitoPerOx-based flow cytometry measuring lipid peroxidation levels as MFI ratios (n = 2 independent experiments). **(Q)** Flow cytometry analysis of mitochondrial mass in LN751 cells treated with PMCi and mitoQ, using MFI ratios (n = 2 independent experiments). **(R)** Schematic model summarizing the mtTIS mechanism described in the manuscript, created using BioRender.com. Statistical analyses, except for **(I),** were performed using one-way ANOVA with Tukey’s multiple comparisons test. Error bars represent mean ± SEM.

To test this hypothesis, we sought to directly assess the role of mitochondrial functionality in PMCi-induced senescence. We generated ρ0 GBM cell lines by depleting mtDNA (fig. S5C) and confirmed the absence of detectable mtDNA in these cells (fig. S5D-F), their inability to execute OXPHOS (fig. S5G) and the absence of ROS production (fig. S5H). Subsequently, we treated these cells with PMCi to ascertain whether the absence of mitochondria affects the induction of senescence by PMCi. In line with our hypothesis, we observed that the treatment failed to induce senescence in these cells. The ρ0 cells did not exhibit characteristic flattening (Fig. 4D and fig. S5I) and failed to produce the SASP molecules associated with PMCi-induced senescence (Fig. 4E). To confirm the contribution of mtDNA depletion to this effect, we conducted a recovery assay to restore mtDNA in the same ρ0 cells (fig. S5C-E). Upon restoration of OXPHOS function (fig. S5G), these cells regained their ability to undergo senescence upon PMCi treatment (Fig. 4D and fig. S5I).

To explore whether the observed mitochondrial dependence is unique to PMCi-induced senescence, we treated ρ0 cells with a high dose of irradiation or a high concentration of palbociclib, two established senescence inducers that are known to generate nuclear DNA damage (Fig. 3A-B and fig. S4A-C). Unlike PMCi, both irradiation and high-dose palbociclib induced senescence regardless of the presence of functional mitochondria, evident from the flattened morphology (Fig. 4F) and the loss of colony potential (Fig. 4G-H and fig. S5J-K). These findings demonstrate that mitochondrial dependence is not a universal feature of all senescence inducing therapies, but distinguishes PMCi-induced senescence and potentially other interventions directly targeting mitochondrial biogenesis or functionality. We propose the term mitochondrial therapy-induced senescence (mtTIS) to describe TIS that is selectively dependent on mitochondria.

### mtTIS requires mtROS-induced mitochondrial damage

After validating the dependence of PMCi-induced senescence on mitochondria we zoomed in on the organelle to see what kind of changes drives mtTIS. Since we saw indications of ROS production by RNAseq (Fig. 3C) and a decreased maximal spare capacity in mitochondrial OXPHOS (Fig. 3M-N and fig. S4I-J), we measured ROS production in PMCi-treated cells. Flow cytometry analysis revealed that PMCi significantly increased mitochondrial ROS (mtROS) in the responsive lines but not in the non-responsive line U251-MG (Fig. 4I and fig. S6A). To assess whether mtROS production is merely a byproduct or a critical factor in senescence induction, we utilized antioxidants to scavenge ROS. Co-treatment with glutathione (GSH) alongside PMCi for 7 days effectively rescued senescence-associated morphology and restored colony formation potential (Fig. 4J-K). However, GSH did not reduce the SA-β-gal signal (fig. S6B). Actually, GSH alone causes SA-β-gal staining without morphological changes of senescence, likely as an artefact of the antioxidant. Notably, the morphological rescue was also evident when GSH was administered only during the first 4 days of treatment (fig. S6C). Similar outcomes were observed with catalase treatment (fig. S6D-F), confirming that ROS scavenging not only mitigates senescent morphology (fig. S6E) but also prevents loss of colony formation potential (fig. S6F). To delineate the involvement of mtROS in this rescue, we employed the mitochondrially-targeted antioxidant mitoquinone (MitoQ) ^32^. MitoQ partially rescued senescence morphology (fig. 4L) and fully restored the colony formation potential of PMCi- treated cells (Fig. 4M) by reducing the amount of mtROS (Fig. 4N).

Given the absence of nuclear DNA damage in PMCi mtTIS and the importance of mtROS production, we hypothesized that the damage might be localized in the mitochondria. We assessed mitochondrial DNA (mtDNA) damage, finding a reduction in mtDNA integrity in senescent cells roughly at level with that incurred by 5 Gy of ionizing radiation, though this was not statistically significant (Fig. 4O). Besides direct DNA damage, ROS can also cause lipid peroxidation, which constitutes an additional mechanism contributing to DNA and organelle damage ^33^. We therefore conducted a mitochondrial lipid peroxidation analysis using flow cytometry. This revealed increased lipid peroxidation in PMCi-induced senescent cells, which was mitigated by the antioxidant MitoQ (Fig. 4P and fig. S6G). As previously shown, MitoQ also rescued the senescent morphology and proliferation potential of PMCi-treated cells (Fig. 4L-M). These findings indicate that mtROS plays a central role in triggering senescence induced by PMCi. Inhibiting mitochondrial biogenesis (Fig. 4A-C), depleting functional mitochondria (Fig 4D-H, and fig. S5H-K), as well as reducing mtROS (Fig. 4J-N,P and fig. S6C-F) all can fully rescue senescence induction, suggesting they function in a linear pathway. Interestingly, MitoQ did not affect the increased mitochondrial mass in senescent cells (Fig. 4Q and fig. S6H), demonstrating that mitochondrial biogenesis precedes mtROS production in PMCi-induced senescence and is not a compensatory response to mtROS generation.

## DISCUSSION

We here identified a context of therapy-induced senescence, in which mitochondria are required and sufficient to drive the senescent state. As such, we labeled this phenomenon mitochondrial therapy-induced senescence, or mtTIS. Prolonged PMCi treatment induced mtTIS by fully shutting down RB phosphorylation, while the continued signaling through the mTOR pathway fuels mitochondrial biosynthesis. This creates an imbalance between cell cycle progression and organelle production. Although the generated mitochondrial network is aberrant, it retains a level of functionality, albeit with limited maximal respiratory capacity. These accumulating dysfunctional mitochondria progressively generate mtROS, leading to mitochondrial damage. In turn, these damage signals are relayed through the cGAS-STING pathway, driving the production of a SASP that induces autocrine and paracrine senescence (Fig. 4R).

PMCi-induced mtTIS distinguishes itself from other senescence contexts in several ways. First and foremost, it occurs in absence of nuclear DNA damage. Classic TIS inducers such as ionizing radiation, chemotherapeutics such as doxorubicin and high concentrations of CDK4/6 inhibitors are widely catalogued to generate nuclear DNA damage ^5^. In fact, the presence of DNA SCARS is generally acknowledged as a classical hallmark of senescence ^34–36^. Setting itself apart, we observe no induction of nuclear DNA damage by PMCi, even though it induces potentially damaging levels of ROS. Instead, the damage is confined to the mitochondria, where both mtDNA and lipid bilayers are affected. Mitochondrial ROS has been long believed to be a source of nuclear DNA damage, but given its reactive and short-lived nature and the distance between mitochondria and nucleus a direct damaging effect is unlikely, as elegantly demonstrated recently by van Soest *et al.* ^37^. A notable exception is micronuclei, which may be in close proximity to mitochondria and can be directly damaged as such ^38^. Of note, we did not observe a significant presence of micronuclei in PMCi responsive cells. Critically, we found that cells lacking mitochondria (ρ0) did not become senescent when exposed to PMCi but these cells were susceptible to nuclear DNA damaging senescence inducers. This clearly established that mitochondria cannot only propagate senescence signaling downstream of nuclear damage signals ^39^, but can actually be the source of the senescence trigger. Previous studies have linked mitochondria to senescent inflammatory signaling through mtDNA leakage via mitochondrial pores, particularly in non-cancer models ^9,40–42^. However, these studies focused predominantly on the role of the mitochondria in SASP induction, finding no involvement in enforcing the durable cell cycle exit. Our findings reveal a clear mitochondrial dependency for both SASP production and cell cycle exit, underscoring mitochondria being capable of driving senescence in absence of nuclear damage. It would be interesting to investigate whether mtTIS depends on mtDNA leakage through mitochondrial pores and whether this contributes to the initiation of senescence signaling. It would also be intriguing to explore whether other organelles could, similar to mitochondria, independently drive senescence under specific conditions.

Second, PMCi-induced mtTIS occurs in cancer cells. Senescence is still predominantly studied in the context of non-transformed cells, such as the MiDAS context described by the late Judith Campisi ^43^. This difference in pre-senescent cell state may be very relevant for PMCi-induced mtTIS, as it is dependent on the imbalance between cell cycle progression and mitochondrial biosynthesis. This imbalance aligns with the concept of ‘toxic overgrowth’ coined by seminal studies from the Saurin, Ly, de Bruin and Neurohr labs ^28–31^. In line with this, an elegant earlier study by Lanz *et al.* demonstrated that cell size is positively correlated with the likelihood of senescence switching ^26^. Intriguingly, so far these studies have not identified which organelle may be responsible for the toxic overgrowth. Our data suggest that mitochondria may be involved in at least some of these contexts. Of note, these studies all concern TIS in cancer cells. This may be critical as mitochondrial biosynthesis is a key driver of toxic overgrowth in mtTIS contexts. The PI3K-AKT-mTOR pathway is a key regulator of mitochondrial biosynthesis and mitochondrial network dynamics ^27^ but also frequently over-activated in cancer, including in GBM ^11^. Cancer cells might therefore be more prone to toxic overgrowth than non-transformed cells. In line with this, we could only observe senescence in cancer cells and not healthy tissues following PMCi exposure in mice.

Third, to the best of our knowledge it has not been previously observed that the SASP cannot just induce paracrine senescence, but is also required to drive senescence in an autocrine fashion in the cells exposed to the original TIS inducer. By driving the accumulation of dysfunctional mitochondria, PMCi indirectly forces the cell to start producing and secreting SASP factors that are sufficient to autocrinely drive the senescent state. Critically, we have conducted rescue experiments at many levels of the proposed mechanism, but could never observe a situation where we found durable cell cycle exit but no SASP. These phenomena always went hand in hand, strongly suggesting that here the SASP is not merely a hallmark associated with senescence and generated by parallel pathways as described before ^9,41,44,45^, but that SASP production precedes senescence switching in PMCi-induced mtTIS and is therefore critical for its induction.

In conclusion, in this study we identify and characterize mtTIS, a previously unappreciated form of senescence for which mitochondria are required. We demonstrate that mitochondria are not just involved in maintaining the senescent state, but can actually be sufficient to drive TIS. Our findings distinguish mtTIS from traditional TIS pathways triggered by nuclear DNA damage, highlighting its relevance for therapies that affect mitochondrial functionality while maintaining nuclear DNA integrity. Finally, this work raises the exciting question whether other cellular organelles may be capable of driving unique senescence contexts.

## MATERIALS AND METHODS

### Serum-cultured cell lines

All cell lines were authenticated using GenePrint 10 STR profiling. LN751 (RRID:CVCL_3964) was kindly provided by Dr. Monika Hegi (CHUV, Lausanne, Switzerland), E98 by Prof. William Leenders, (RUMC, Nijmegen, The Netherlands), U251-MG (RRID:CVCL_0021) by Dr Martha Chekenya and Dr. Rolf Bjerkvig (University of Bergen, Norway), LN428 (RRID:CVCL_3959) and T98G (RRID:CVCL_0556) by Dr. Conchita Vens (NKI, Amsterdam, The Netherlands). U87-MG (RRID:CVCL_0022), U118-MG (RRID:CVCL_0633) and HS683 (RRID:CVCL_0844) were purchased from ATCC (Manassas, VA). All human GBM cell lines were cultured and maintained in complete Minimal Essential Medium (cMEM): MEM supplemented with 10% Fetal Bovine Serum (FBS), 1% L-glutamine, 1% MEM-Vitamins, 1% Non-Essential Amino Acids, 1% Penicillin-Streptomycin and 1% Sodium Pyruvate (all purchased from Invitrogen, Grand Island, NY) and grown in standard conditions (37 °C and 5% CO_2_).

### Glioblastoma stem cell lines

All GSC cultures were authenticated using GenePrint 10 STR analysis. Human glioblastoma stem cell lines (GSCs) U3086-MG (RRID:CVCL_IR96), U3065-MG (RRID:CVCL_IR87) and U3013-MG (RRID:CVCL_IR61) were obtained from the Human Glioma Cell Culture resource^46^. The GBM8 GSC line was kindly provided by dr. Bakhos Tannous (Massachusetts General Hospital, Boston, MA). The murine glioma stem cell lines GSC578 (*Egfr^viii^;Cdkn2a/b^-/-^;Pten^-/-^*) and GSC457 (*Kras^v12^;Cdkn2a^-/-^;Tp53^-/^*^-^) were isolated from spontaneous glioma models generated through the intracranial injection of lentiviral Cre vectors into LoxP-conditional mice, as described previously ^47–49^. All GSCs were cultured in medium consisting of 1:1 neurobasal medium and DMEM GlutaMAX high glucose-Ham’s F12 nutrient mix, supplemented with 2% B27 without vitamin A, 1% Penicillin/Streptomycin, 10 ng/mL epidermal frowth factor (EGF) and 10 ng/mL fibroblast growth factor (FGF) (all originated from Thermo Fisher Scientific, Waltham, MA). Since we have demonstrated that antioxidants rescue senescence we decided to treat the stem cells in antioxidant free medium. Therefore, in each experiment, GSC lines were cultured in MHM, a B27-free stem cell medium consisting of DMEM/F12 supplemented with 0.6% glucose, 0.15% NaHCO_3_, 22 mM HEPES, 2 mM L-glutamine, 100 units/mL penicillin, 100 μg/mL streptomycin, 10 ng/mL EGF, 10 ng/mL FGF, 0.1 mg/mL transferrin, 10 μg/mL putrescine, 25 μg/mL insulin, 50 ng/mL sodium selenite, and 60 ng/mL progesterone (all Sigma-Aldrich, St. Louis, MO). To culture the cells adherently, culture dishes were coated with 1.5 mg/mL poly-ornithine (Sigma-Aldrich) for at least 3 hours at 37 °C, after which the plates were washed, and 1 μg/mL laminin (Sigma-Aldrich) was added for an hour at 37 °C.

### Compounds

Buparlisib was obtained from Syncom (Groningen, Netherlands); mirdametinib from LC laboratories (Woburn, MA); palbociclib and abemaciclib from MedKoo Biosciences (Morrisville, NC); MK2206 from Active Biochemicals (Wan Chai, Hong Kong) and samatolisib, trametinib, AZD8055, H-151, BAI1 and mitoquionone from MedChemExpress (Sollentuna, Sweden). All drugs were dissolved in dimethylsulfoxide (DMSO; Sigma-Aldrich). Prior to each experiment, drugs were further diluted in the appropriate culture medium.

### Ionizing radiation

Cells were seeded in 6-well plates at a density suitable for subsequent analysis and allowed to adhere overnight under standard conditions (37°C, 5% CO). The following day, cells were irradiated using a ¹³ Cs irradiator (Gammacell-1000 Unit). The irradiation dose was calibrated based on experimental requirements. After irradiation, the cells were returned to standard culture conditions and maintained for 7 days before further analysis.

### Simple Western Analysis

Cells were harvested using a cell scraper (Corning Inc., Corning, NY), washed with ice-cold PBS, and centrifuged at 800 rcf for 5 minutes. The cell pellet was lysed in ice-cold lysis buffer (0.1% Triton X-100, 20 mM Tris-HCl [pH 7.6], 100 mM NaCl) supplemented with 2 mM sodium fluoride, 1 mM sodium orthovanadate, 1 mM sodium pyrophosphate decaldehyde, 2.5 mM β-glycerophosphate, 1× protease inhibitor cocktail, 1 mM phenylmethylsulfonyl fluoride (PMSF), and 1 mM dithiothreitol (DTT). Lysis was conducted on ice for 30 minutes with intermittent vortexing. The lysates were centrifuged at 16,100 rcf for 15 minutes at 4 °C and protein concentration was quantified using the DC™ Protein Assay kit (Bio-Rad, Hercules, CA).

For Simple Western analysis, lysates were prepared per manufacturer’s instructions. Automated protein separation and immunodetection were carried out using the Jess Simple Western system (ProteinSimple, San Jose, CA). Antibodies used (Supplementary Table S1) included phospho-Akt, phospho-Erk, phospho-Rb and FOXM1. Data were analyzed using Compass software (ProteinSimple).

### Endpoint proliferation assays

Cell proliferation assays were used to assess the effect of PMCi on cell proliferation and morphological changes. Cell densities were optimized for each cell line. Cells were seeded in each well of a 24-well plate with either 500 µl cMEM for serum-cultured cell lines or MHM for GSCs. The next day, drugs were added to each well of the plates to a final volume of 1 mL. For the PMCi the final drug concentrations in each well ranged from 100 nM to 500 nM for buparlisib, 1 nM to 10 nM for mirdametinib, and 10 nM to 100 nM for palbociclib. A minimum of two independent experiments were performed for each inhibitor. For the rest of the compounds the final concentrations were 2 mM of glutathione (GSH), 2000 U/mL catalase, 2 mM n-acetyl cysteine (NAC), 100 nM mitoquinone (mitoQ), 3 μM for MK2206 and 10 μM for GSK690693. The control consisted of either cMEM or MHM without drugs. Plates were incubated for seven days for senescence induction. In that time ∼90% confluency was reached in control wells. After incubation, cells were stained and fixated with glutaraldehyde (6% v/v) and crystal violet (0.5% w/v). Images of the plates were taken with the ChemiDocTM MP system (Bio-Rad) using the Image Lab Software. Cell density was analyzed with FIJI ^50^ by applying the Colony Area plugin ^51^. Photos of each condition were taken by applying bright-field microscopy using a CCD system (Carl Zeiss Microscopy LLC, Thornwood, NY).

### Incycute experiments

Cells were seeded in 24-well plates at a density optimized for each cell type and allowed to adhere overnight under standard culture conditions (37°C, 5% CO). The following day, the plates were transferred to the Incucyte® S3 system (Sartorius) for live-cell imaging and confluency monitoring.

The Incucyte® S3 system was configured to capture phase-contrast images using a 4x objective lens, with images taken every 1 hour. The standard confluency analysis settings in the Incucyte® software were used to monitor cell growth and confluency over several days. The system automatically analyzed the images to calculate the confluency of each well, providing a quantitative assessment of cell proliferation over time. Data were collected and analyzed using the Incucyte® S3 software, with results expressed as mean confluency percentages normalized to the control wells.

### *In vitro* senescence-associated **β**-galactosidase assay

To further validate the induction of senescence by PMCi, cell lines were stained using the senescence-associated ß-galactosidase (SA-β-gal) staining assay according to the staining protocol described by Itahana *et al*. ^52^. Images were taken with bright-field microscopy using a CCD system (Carl Zeiss Microscopy). Two independent experiments were performed for each cell line.

### Cytokine arrays

Cells were seeded in 6-well plates and allowed to adhere overnight under standard conditions of 5% CO_2_ and 37 °C. Subsequently, cells were either left untreated in cMEM or submitted to the different treated conditions for 7 days. On the 7-day time point, cells were washed with PBS, followed by the addition of 1 ml cMEM to each well. After 24 hours, the conditioned medium was collected and stored at −20°C until further analysis. Conditioned medium samples were subjected to analysis using the Human XL Cytokine Array Kit (R&D Systems, Inc., Minneapolis, MN)per manufacturer’s protocol. This assay facilitated the simultaneous detection of multiple cytokines. To account for potential variations in cell proliferation induced by the treatment, both treated and control cells were counted immediately following conditioned medium collection. The obtained results were subsequently normalized to the respective cell numbers.

### Conditioned medium

Cells were cultured for 7 days with PMCi treatment to induce senescence, confirmed by bright-field microscopy. Following the treatment period, the drug-containing medium was replaced with cMEM. On day 10 or day 14 from the start of the treatment, the conditioned medium (CM) was collected and filtered using a 0.22 µm Millex-GS Syringe Filter (Millipore Sigma, Darmstadt, Germany). The filtered CM was then applied to freshly seeded cells to assess the effects of the SASP.

As a control, cells were treated with CM collected from untreated cells cultured for 7 days in cMEM to account for any growth factor depletion that might occur during CM generation (“control” CM). After a 7-day incubation with either senescent or control CM, cells were stained and fixed using a solution of 6% glutaraldehyde (v/v) and 0.5% crystal violet (w/v) to assess morphological changes or stained for SA-β-gal activity to determine senescence induction. Imaging was performed using the Zeiss Observer Z1 microscope at 20x magnification with phase-contrast microscopy. All experiments were independently repeated three times.

### Quantification of mitochondrial ROS, lipid peroxidation, and mass

Specific fluorescent indicators were employed to assess changes in mtROS (MitoSOX Red; Thermo Fisher Scientific), mitochondrial mass (MitoTracker Green FM and MitoTracker Deep Red; (Invitrogen, Waltham, MA)), and mitochondrial lipid peroxidation (MitoPerOx; Abcam Ltd, Cambridge, UK). Cells were treated with the respective indicators and incubated for 30 minutes. Following the incubation period, cells were trypsinized, stained with DAPI, and subjected to analysis using an Attune NxT Flow Cytometer (Thermo Fisher Scientific). Flowjo Single Cell Analysis Software v10 (Flowjo LLC, Ashland, OR) was employed for gating and analysis. The population of interest was identified through consecutive gating steps, utilizing 1. SSC-Area vs. FSC-Area (cells ecxluding debris), 2. SSC-Height vs. SSC-Area (single cells), 3. VL1-Height (DAPI) vs. BL1-Area, BL2-Area or RL1-Area (live cells depending on the dye to be analyzed). Median Fluorescence Intensity (MFI) values were calculated from the final live cell gate. Of all conditions at least two independent biological replicates were analyzed per experiment and at least two independent experiments were performed. Background fluorescence was subtracted using the average value from two independent biological samples of unstained cells (fluorescence minus-one (FMO) controls) for each condition, resulting in an MFI value for each stained sample. MFI values from every stained sample was then normalized to the average MFI value of the control condition, resulting in an MFI ratio.

### Comet assay

For the initial preparation, cells were washed with Hanks’ Balanced Salt Solution (HBSS) and dissociated with trypsin-EDTA (Thermo Fisher Scientific, Waltham, MA). After trypsinization, the cells were resuspended in cMEM. For the comet assay, 5000 cells were added to PBS at 4°C, followed by the addition of 1% agarose. The mixture was vortexed rapidly to ensure uniform cell distribution and immediately pipetted onto Epredia™ SuperFrost Plus™ Adhesion slides. The comet slides were carefully placed in a humid chamber to avoid evaporation and stored on ice to allow the agarose to set properly. Afterwards, the slides were submerged in the lysis solution (pH=10, 2.5 M NaCl, 0.1 M Na_2_EDTA, 10 mM Tris, 1% Sarkosyl, 1% Triton-X100) and stored at 4°C to ensure optimal lysis. After lysis, the slides were washed with alkaline (pH > 13, 200 mM NaOH, 2 mM Na_2_EDTA) or neutral (pH = 8, 90 mM TrisHCL, 90 mM Boric Acid, 2 mM Na_2_EDTA) running buffer at room temperature. The slides were then submerged in the electrophoresis chamber, and electrophoresis was conducted for 25 minutes at 20 V in 4 °C. Post-electrophoresis, slides were fixed with 70% ethanol for 30 minutes at room temperature and then dried at 37 °C for 15 minutes. Staining was performed for 20 minutes using a final concentration 2.5 µg/mL propidium iodide (PI) in distilled water. Slide analysis was conducted using the Zeiss Observer Z1 microscope, using a 20x Phase 2 objective with the 31 Alexa Fluor 568 filter. Comet analysis was carried out using OpenComet v1.3.1 software. The software enabled the automated measurement of various comet parameters, including the Olive Tail Moment, which was used to assess the extent of DNA damage.

### γH2AX fluorescence microscopy

Cells were grown on glass coverslips and pre-extracted with cold PBS containing 0.2% Triton-X100 on ice. After the were fixed using 3.7% formaldehyde, supplemented with 0.2% Triton-X100, at 4 °C. Coverslips were placed on parafilm within a humidified chamber. Cells were incubated with blocking buffer containing 2% Bovine Serum Albumin (BSA) in PBS and 0.1% Triton. After removing the blocking buffer we incubated them with primary anti-phospho-γH2AX antibody (CST9718, Cell Signaling Technology, Danvers, MA) in 1:1000 dilution. The coverslips were then incubated overnight at 4 °C. Post-incubation, coverslips were washed twice with PBS containing 0.5% Tween 20 (PBS-T) and the secondary goat-anti-rabbit-AF488 (AB150077, Abcam) antibody in 1:500 dilution incubated for 2 hours at room temperature. Finally, the coverslips were washed twice with PBS-T and then stained with ProLong™ Diamond Antifade Mountant with DAPI (Thermo Fisher Scientific). The foci photos were taken with the Zeiss Observer Z1 microscope, using a 20x Phase 2 objective. The foci were analyzed using the *Foci_Analyzer_1_3.ijm* script in Fiji (ImageJ) found in https://github.com/BioImaging-NKI/Foci-analyzer.

### Mitochondrial immunofluorescence microscopy

Mitochondrial membranes were stained using MitoTracker Green FM (Invitrogen, Waltham, MA) to analyze the mitochondrial network structure. The dye was applied at a concentration of 200 nM for 45 minutes, following the manufacturer’s instructions. After staining, cells were seeded in a 35 mm µ-Dish (ibidi, Gräfelfing, Germany) and allowed to grow for 48 hours under standard culture conditions. Imaging was conducted using an Andor Dragonfly system (Andor, Belfast, Northern Ireland), installed on an inverted Leica DMI8 microscope equipped with a 40x oil immersion objective.

### Creation of **ρ**0 cells

Mitochondrial DNA (mtDNA) depletion, or the creation of ρ0 cells, was performed according to previously published literature ^53^, using 100 ng/mL ethidium bromide (EtBr) for a duration of 1 month. Depletion of mtDNA was confirmed using SYBR Gold dye (Thermo Fisher Scientific), which labels mitochondrial nucleoids, and MitoTracker Red FM (Invitrogen, Waltham, MA), which selectively stains mitochondria in live cells. The dyes were applied according to the manufacturer’s instructions, with SYBR Gold at a 1:10000 dilution and MitoTracker Red FM at a concentration of 200 nM for 45 minutes. Following staining, cells were seeded in a 35 mm µ-Dish (ibidi, Gräfelfing, Germany) and imaged 48 hours later. Imaging was performed using either a TCS SP5 Confocal microscope (Leica Camera AG, Wetzlar, Germany) with a 63x oil immersion objective or an Andor Dragonfly system (Andor, Belfast, Northern Ireland) installed on an inverted Leica DMI8 microscope with a 40x oil immersion objective.

### mtDNA copy number quantification

DNA isolation was performed utilizing the QIAamp DNA Mini Kit (Qiagen, Hilden, Germany). Subsequently, mitochondrial DNA (mtDNA) was selectively targeted through the design of custom primers (see Supplementary Table S3 for primer details) and optimizing melting curves and validating optimal amplification efficiency. To assess the relative mtDNA content compared to nuclear DNA, the ΔCt value was determined by subtracting the average cycle threshold (Ct) value of the mitochondrial target from that of the nuclear target. Subsequently, the ΔΔCt values were calculated by subtracting the control ΔCt from the corresponding values of the experimental conditions. The 2^ΔΔCt formula was applied to obtain the relative quantification of mtDNA content compared to the control.

### mtDNA Damage Analysis

To quantify mtDNA damage, isolated DNA was analyzed using the Human Real-Time PCR Mitochondrial DNA Damage Analysis Kit (Detroit R&D, Detroit, MI) according to the manufacturer’s instructions. mtDNA damage was assessed by comparing the amplification efficiency of long and short mtDNA fragments. Higher Ct values for the longer 8.8 kb fragment indicate greater mtDNA damage, as a high level of this product corresponds to lower levels of mtDNA damage.A standard curve was generated using the threshold cycle (Ct) values and the DNA concentration (log scale) of each 8.8 kb standard. Linear regression analysis of this curve was used to determine the DNA concentration of each sample based on its Ct value obtained through PCR. The formula 10^mtDNA concentration was applied to calculate relative mtDNA integrity. The values were then normalized to express fold change relative to control samples, providing a measure of mtDNA damage..

### Oxygen consumption rate measurement

To assess the oxygen consumption rate (OCR), cells were seeded in a Seahorse XFe24 cell culture microplate (Agilent) at a density of 60,000-70,000 cells per well, optimized for the cell type, and incubated overnight at 37°C with 5% CO to allow for cell adherence. On the day of the assay, the cell culture medium was replaced with DMEM supplemented with 10 mM glucose, 1 mM pyruvate, and 2 mM glutamine, adjusted to pH 7.4) (all originated from Thermo Fisher Scientific, Waltham, MA). The plate was incubated at 37°C in a non-CO incubator for 10 minutes to allow for temperature and pH equilibration. Compounds for the mitochondrial stress test, including oligomycin (1 µM, Sigma-Aldrich), FCCP (1 µM, Sigma-Aldrich), and a mix of rotenone and antimycin A (1 µM each, Sigma-Aldrich), all dissolved in dimethylsulfoxide (DMSO; Sigma-Aldrich), were loaded into the appropriate injection ports of the hydrated sensor cartridge. The cell plate and loaded sensor cartridge were then placed into the Seahorse XFe24 Analyzer, and the assay was programmed using the Seahorse Wave software. The assay protocol included sensor calibration, followed by four baseline OCR measurements (4 minutes mix, 4 minutes measure) before sequential injections of oligomycin, FCCP, and rotenone/antimycin A, with three measurement cycles following each injection. Data were collected and analyzed using the Seahorse Wave software to determine parameters such as basal respiration, ATP production, maximal respiration, and spare respiratory capacity. Following the assay, cells were rinsed with PBS, and cell counts were measured to normalize OCR values. This normalization ensured that OCR data accurately reflected cellular metabolic activity per cell..

### Clonal outgrowth capacity assessment

To assess the colony formation ability of cells were seeded in T75 and incubated overnight under normal conditions (37 °C and 5% CO_2_). The following day, flasks were treated with cMEM, PMCi, inhibitor or PMCi + inhibitor. Two weeks post-treatment, cells were washed with HBSS, trypsinized, and centrifuged. The resulting cell pellets were diluted in 1 mL of PBS. Cells from each condition were sorted into three 96-well plates containing 100 μL cMEM to create triplicates. A FACSAria™ Fusion flow cytometer (BD Biosciences, Franklin Lakes, NJ) was utilized for cell sorting. In experiments involving ρ0 cells, a modified procedure was followed. After treating the cells with different compounds, they were replated at a low density of 100-200 cells per well in a 6-well plate. Two weeks post-sorting or replating, plates were stained and fixed using a solution containing glutaraldehyde (6% v/v) and crystal violet (0.5% w/v). The quantification of colonies was manually performed by two independent blinded observers using bright-field light emission microscopy. Positivity for colony formation was assigned to colonies containing more than three cells.

### Electron microscopy

Adherent cells were fixed in a solution containing 2% paraformaldehyde and 2.5% glutaraldehyde in 0.1 M sodium cacodylate buffer (pH 7.2) for 1 hour at room temperature, followed by overnight fixation at 4°C. After fixation, specimens were thoroughly washed to remove any residual fixatives. Post-fixation staining was performed using 1% osmium tetroxide for 2 hours at room temperature. Subsequently, the specimens underwent en-bloc staining with 1% uranyl acetate for 1 hour at 37°C, followed by additional washing steps to eliminate excess staining agents. The specimens were then subjected to a graded ethanol dehydration series, consisting of 50% ethanol, two rounds of 70% ethanol, 90% ethanol, and three rounds of 100% ethanol. Following dehydration, BEEM capsules filled with a plastic mix were inverted directly onto the cells while they remained submerged in 100% ethanol. The plastic mix was polymerized for 48 hours at 60°C, ensuring that the cells remained in their original orientation, facilitating direct comparison between different conditions. TEM images were acquired using a JEM-F200 Multi-purpose Electron Microscope (JEOL Ltd., Tokyo, Japan).

### Animals

Animal housing and studies were conducted in compliance with national regulations and institutional guidelines and received approval from the institutional animal experimental committee. The experiments were conducted under license AVD301002016407 with working protocol number 1.1.8523. Animals were housed in a temperature-controlled environment with a 12-hour light/dark cycle and were provided with *ad libitum* access to food and water.

### Orthotopic xenograft studies

Orthotopic GBM tumors were induced in *Abcb1a/b;Abcg2^-/-^*mice by stereotactic injection of E98 cells expressing firefly luciferase ^54^. At an injection speed of 1 µL/min, 200,000 cells were injected in 2 µL HBSS (Invitrogen) 2 mm lateral, 1 mm anterior and 3 mm ventral to the bregma. Tumor growth was monitored using bioluminescence imaging by injecting D-luciferin (150 mg/kg i.p.; Promega, Madison, WI) and subsequently imaging mice on an IVIS Spectrum Imaging System (PerkinElmer, Waltham, MA). Survival was monitored daily and the humane endpoint was defined as body weight loss exceeding 20%. Male and female tumor-bearing mice were treated for 14 days by supplying acidified vehicle drinking water (HCl, pH4) consisting of 2% w/v (2-Hydroxypropyl)-β-cyclodextrin and 10% w/v sucrose (both Sigma-Aldrich) or vehicle water containing 60 μg/ml buparlisib, 30 μg/ml mirdametinib and 150 μg/ml palbociclib.

### Pharmacokinetic studies

Pharmacokinetic studies were conducted using *Abcg2^-/-^;Abcb1a/b^-/-^*FVB nude mice as previously described ^55^. Plasma samples were subjected to liquid-liquid extraction with diethyl ether. The residues were reconstituted in acetonitrile: water (30:70 v/v) and analyzed using an LC-MS/MS set-up comprising an Ultimate 3000 LC System (Dionex, Sunnyvale, CA) and an API 3000 mass spectrometer (AB Sciex, Framingham, MA). Separation was carried out on a ZORBAX Extend-C18 column (Agilent Technologies, Santa Clara, CA), with a mobile phase gradient from 30% B to 95% B over 5 minutes, maintained for 3 minutes, followed by re-equilibration at 30% B. The data were analyzed using Analyst 1.5.1 software (AB Sciex).

### Histology and immunohistochemistry

E98 tumor-bearing brain and healthy skin were harvested from *Abcb1a/b;Abcg2^-/-^* mice at the end of drinking water treatment (14 days). Tissues were embedded in Tissue-Tek OCT Compound (Sakura Finetek, Alphen aan den Rijn, the Netherlands) and cut in 8 µm sections using a cryostat. Sections were stained for SA-β-gal activity using the CS0030 histochemical staining kit and counterstained using nuclear fast red (both Sigma-Aldrich) with or without anti-vimentin staining (1:200, CST5741, Cell Signaling Technology) as indicated.

### Blood parameter analysis

Whole blood was collected from PMCi-treated mice by cardiac puncture immediately after sacrificing using CO_2_ at the final day of treatment. Heparin was added to prevent clotting and whole blood samples were analyzed using a DxH500 Hematology Analyzer (Beckman Coulter, Brea, CA).

### RNAseq

Total RNA was extracted from LN751 and U251-MG cells with the RNeasy mini kit (QIAGEN), including a column DNase digestion (QIAGEN) according to the manufacturer’s instruction. Quality and quantity of total RNA was assessed by the 2100 Bioanalyzer using a Nano chip (Agilent, Santa Clara, CA). Strand-specific libraries were generated using the TruSeq Stranded mRNA samples preparation kit (Illumina Inc., San Diego, CA) according to the manufacturer’s instructions. Briefly, polyadenylated RNA from intact total RNA was purified using oligo-dT beads. Following purification, the RNA was fragmented, random primed, and reverse transcribed using SuperScript II Reverse Transcriptase (Invitrogen, Waltham, MA) with the addition of Actinomycin D. Second strand synthesis was performed using Polymerase I and RNaseH with replacement of dTTP for dUTP. The generated cDNA fragments were 30 end adenylated and ligated to Illumina Paired-end sequencing adapters and subsequently amplified by 12 cycles of PCR. The libraries were analyzed on a 2100 Bioanalyzer using a 7500 chip (Agilent), diluted, and pooled equimolar into a multiplex sequencing pool. The libraries were sequenced with single-end 65bp reads on a HiSeq 2500 using V4 chemistry (Illumina Inc.).

For the analysis of RNAseq data, we followed two approaches. First, for differential expression and GSEA analyses, gencode v34 protein-coding transcripts were quantified using salmon (v1.9.0) ^56^ followed by differential analysis and GSEA using the DESEQ2 ^57^ and clusterprofiler ^58^ packages in the R environment. For the GSEA, gene ranking was performed using the log2FC x min(-log10(padj),3) measure and gene sets were retrieved from MSigDb v7.0 ^59^. To analyze cell cycle phase-specific expression patterns, we performed an ssGSEA analysis using the marker genes from ^60^. In this analysis, after calculating the ssGSEA scores for each sample and for each phase-specific gene set, we calculated the average ssGSEA score for each group (untreated CTRL, 7d, 14d & 21d treatment), and plotted the delta ssGSEA scores for each cell-cycle phase and group pair. This approached allowed us to visualize the relative differences between phase-specific expression patterns. In our second approach, senescence scores were calculated using the SENCAN classifier described in Jochems *et al.* ^13^, by following the same quantification pipeline proposed in this paper. For this approach, transcript levels were quantified using kallisto (v0.46) and summed to gene level. As reference transcriptome, Gencode v34 was used, where we filtered out transcripts from genes annotated as unexpressed pseudogenes as well as transcripts with a transcript support level of 5 unless the transcript has a consensus coding sequence identifier. The gene expression levels were normalized between samples with edgeR (v3.26.8) using the trimmed mean of M-values with singleton pairing. The (untransformed) gene expression counts from kallisto were used directly as input for the SENCAN classifier. MAPK pathway activity scores were calculated exactly as described before ^61^, using the log2 counts per million of the 10 signature genes as input. The average expression of mitochondrial genes was estimated by summing the non-transformed counts per million for the 15 mitochondrial-encoded genes, followed by log2 transformation.

The RNAseq dataset generated and analyzed in this study is available through the NCBI Gene Expression Omnibus database under accession ID GSE277882.

### Proteomics

LN751, E98 and U251-MG cells were lysed, reduced and alkylated in heated guanidine (GuHCl) lysis buffer as described by Jersie-Christensen *et al.* ^62^. Protein concentrations were determined with a Pierce Coomassie (Bradford) Protein Assay Kit (Thermo Fischer Scientific), according to the manufacturer’s instructions. After dilution to <2M GuHCl, aliquots comprising 1100 µg of protein were digested twice (4h and overnight) with trypsin (TPCK-treated, Sigma-Aldrich) at 37°C, enzyme/substrate ratio 1:50. Digestion was quenched by the addition of formic acid (final concentration 5%), after which peptides were desalted on a Sep-Pak 3cc tC18 cartridge (Waters Corporation, Milford, MA), vacuum-dried and stored at −80 °C until further processing.

For proteome analysis, triplicate equal digest amounts of control, 1h-treated and PMCi-treated cells were labeled for each cell line with TMT-126, -127N/C, -128N/C, -129N/C and -130N/C isobaric tags (Thermo Fischer Scientific) according to the manufacturer’s instructions. After checking for labeling completeness, digests were mixed in a 1:1 fashion in such a way that for each cell line a pool of 9 samples (three replicates 0h, 1h and PMCi) was created. Each cell line pool was subjected to basic reversed-phase (HpH-RP) high-performance liquid chromatography for offline peptide fractionation as described previously ^63^, with a slight modification in gradient steepness to account for increased peptide hydrophobicity stemming from TMT labeling. Collected fractions were concatenated to 12 fractions, vacuum dried and aliquots comprising ∼1 µg of peptide of each concatenated fraction were analyzed by nanoLC-MS/MS on an Q Exactive HF-X Hybrid Quadrupole-Orbitrap Mass Spectrometer, equipped with an EASY-nLC 1200 system (Thermo Fischer Scientific). Samples were directly loaded onto the analytical column (ReproSil-Pur 120 C18-AQ, 2.4 μm, 75 μm × 500 mm, packed in-house with integrated emitter). Solvent A was 0.1% formic acid/water and solvent B was 0.1% formic acid/80% acetonitrile. Samples were eluted from the analytical column at a constant flow of 250 nl/min in a 124-min gradient containing a linear increase from 9-30% solvent B. Nanospray was achieved using the Nanospray Flex Ion Source (Thermo Fischer Scientific) in liquid junction setup with spray voltage set to 2.1 kV. The mass spectrometer was operated in Top 10 data-dependent acquisition (DDA) mode, with full MS scans being collected in the Orbitrap analyzer with 60,000 resolution at m/z 200 over a 375-1500 m/z range. Default charge state was set to 2+; the AGC target was set to 3e6; maximum injection time was 25 ms and dynamic exclusion time was set to 17.5 s. For ddMS2, a normalized HCD collision energy of 28% was applied to precursors with a 2+ to 6+ charge state meeting a 1e5 intensity threshold filter. Fixed first mass was set to m/z 110 because of TMT; AGC target was 1.00e4; precursors were isolated in the Quadrupole analyzer with a 1.0 m/z isolation window and MS2 spectra were acquired at 45,000 resolution in the Orbitrap.

Proteome data (RAW files) were analyzed by Proteome Discoverer (version PD 2.5.0.400, Thermo Fischer Scientific) using standard settings. MS/MS spectra were searched against the human Swissprot database (20375 entries, release 2020_09) complemented with a list of common contaminants and concatenated with the reversed version of all sequences, using Sequest HT. The maximum allowed precursor mass tolerance was 50 ppm and 0.06 Da for fragment ion masses. Trypsin was chosen as cleavage specificity allowing two miscleavages. Carbamidomethylation (C), TMT6plex (K) and TMT6plex (N-term) were set as a fixed modifications, whereas Ac (Protein N-terminus) and oxidation (M) were selected as variable modifications. False discovery rates for peptide and protein identification were set to 1% and as additional filter Sequest HT XCorr>1 was applied. TMT quantification was performed using standard settings. The most confident centroid integration method was selected within the “Reporter Ions Quantifier” processing node, whereas reporter abundances were based on S/N (“Automatic”) and the co-Isolation Threshold was set to 30 within the “Reporter Ions Quantifier” consensus node. For each cell line, the PD output file containing 9 TMT channel protein abundances were loaded into Perseus (version 1.6.14.0) ^64^. Abundances were Log2-transformed and proteins were filtered for valid values (abundances determined) in at least all 3 replicates of one condition.

The mass spectrometry proteomics data have been deposited to the ProteomeXchange Consortium via the PRIDE ^62^ partner repository with the dataset identifier PXD056691.

### Statistical analysis

Statistical analyses were conducted using GraphPad Prism software. Specific details for each analysis are provided in the corresponding figure legends. Bar graphs comparing two conditions (treated vs. untreated) were analyzed using unpaired Student’s t-tests. When comparing multiple treatment conditions in one cell line, one-way ANOVA followed by Tukey’s multiple comparisons test was employed. To examine the effects of two independent variables on a dependent variable, we applied two-way ANOVA with Šidák’s test. For mouse survival curves, the log-rank (Mantel-Cox) test was applied to compare survival distributions between groups. For proteomic and transcriptomic data, specialized statistical methods were employed as described above in the corresponding method sections.

## Acknowledgments

The authors thank Piotr Waranecki for conducting the STR analyses. The authors thank Marjolijn Mertz, Amalie Dick and Lenny Brocks from the NKI BioImaging Core Facility for their assistance with image processing and analysis. The authors thank Lex de Vrije from the NKI Animal Pathology Core Facility for assistance with (immuno-)histochemistry. The authors thank Frank van Diepen and Martijn van Baalen from the NKI Flow Cytometry Core Facility for assistance in setting up flow cytometry assays.

## Funding

Dutch Cancer Society (KWF) grant 13148 (OvT, MCdG).

Stichting STOPHersentumoren (OvT).

Stichting Aniek Hendriksz Second to None (OvT).

## Author contributions

Conceptualization: CB, OvT, MCdG

Methodology: CB, FA, IMP, MGN, BT, FJ, OB, HJ, LEK

Investigation: CB, FA, IMP, MGN, BT, RL, KP, DMF, AT, KR, OB, HJ, OvT, MCdG

Visualization: CB, OvT, MCdG Funding acquisition: OvT, MCdG

Project administration: CB, OvT, MCdG

Supervision: JHB, WJF, JS, RB, OvT, MCdG

Writing—original draft: CB, OvT, MCdG

Writing—review & editing: CB, FA, RL, FJ, LEK, WJF, JS, RB, OvT, MCdG

