## Supplementary Materials for "Mitochondrial damage triggers therapy-induced senescence"

Chrysiida Baltira *et al.*

*Corresponding authors.
Olaf van Tellingen
Mark de Gooijer

**This file includes:**

Figs. S1 to S6
Tables S1 to S3

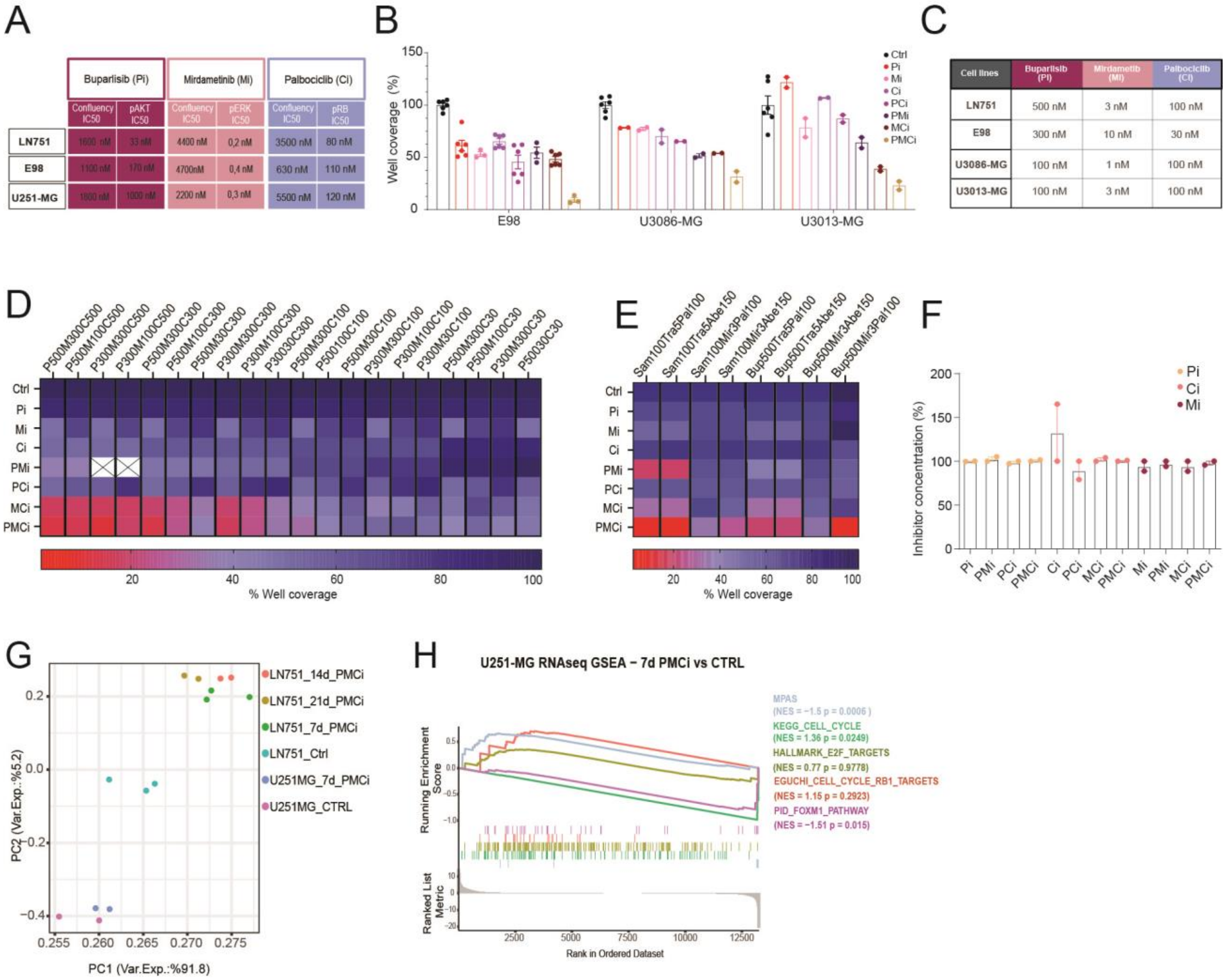


**Fig. S1**. **Triple combination of PMCi target compounds has superior efficiency in inhibiting cell proliferation compared to single or double dosages. (A)**Table summarizing IC50 values of proliferation (Incucyte) and target inhibition (Simple Western) for LN751, E98, and U251-MG **(B)** Proliferation of E98, U3086-MG and U3013-MG treated with single, double, and triple combinations of PMCi after optimization concentrations for each cell line. Minimum n = 2 independent experiments. **(C)** Optimal PMCi concentrations for the most effective inhibition of cell proliferation in triple combinations for each cell line. These concentrations correspond to those used in Figure 1F-G. **(D)** Heat map depicting cell proliferation in U251 cells after 7 days of treatment with single, double, or triple combinations of PMCi compounds. The experiment was conducted in two independent replicates. Compounds: P = Buparlisib, M = Mirdametinib, C = Palbociclib. **(E)** Heat map showing cell proliferation in LN751 cells after 7 days of treatment with single, double, or triple combinations of PMCi compounds. Compounds: Bup = Buparlisib, Mir = Mirdametinib, Abe = Abemaciclib, Pal = Palbociclib, Tra = Trametinib, Sam = Samotolisib. **(F)** Bar graph representing the stability of the PMCi drugs diluted in cMEM and added to GBM cells. Stability was measured by LC-MS /MS after 7 days of treatment. **(G)** Principal Component Analysis (PCA) illustrating the clustering of LN751 and U251 cell lines based on their responses to PMCi treatment along Principal Component 1 (PC1) and Principal Component 2 (PC2). **(H)** GSEA comparing cells treated with PMCi for 7 days to control cells in the non-responsive U251-MG cell line. The plot shows normalized enrichment scores (NES) for various gene sets, with different colors indicating distinct gene sets and the corresponding p values. Error bars represent mean ± SEM.

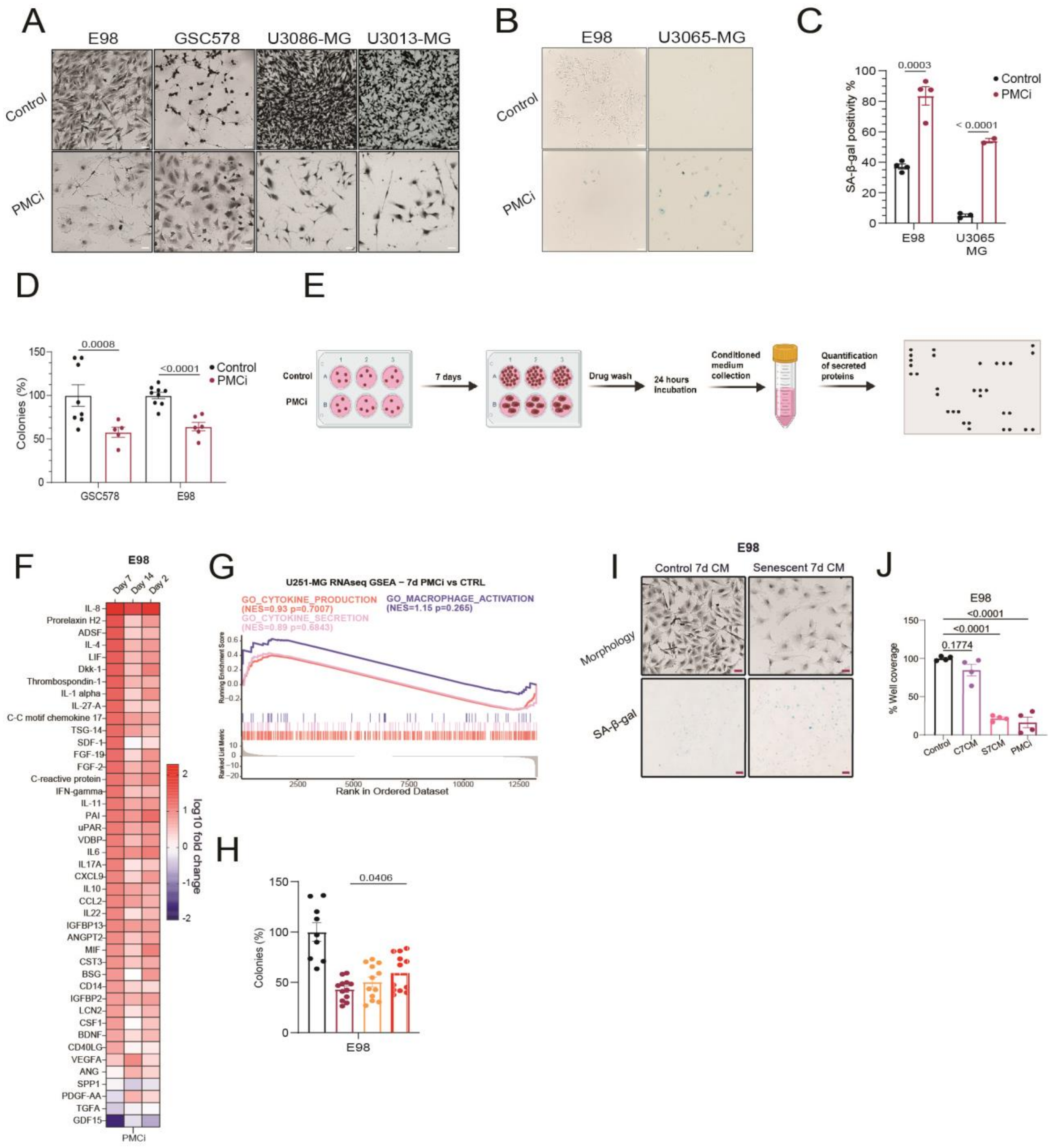

**Fig. S2. PMCi senescent cells produce a functional SASP with common senescent factors. (A)** Representative photographs of the morphological changes after 7 days of PMCi treatment (scale bar is 50uM) in four PMCi-senescence responsive lines **(B)** Representative SA-β- galactosidase staining photographs (scale bar is 100uM) in two responsive lines and **(C)** Quantification in the same cell lines (n = 2 independent experiments). Separate experiments per cell line were analyzed with unpaired Student’s t-test. **(D)** Colony formation assay quantification in two responsive GBM lines. Minimum n=2 independent experiments. Separate experiments per cell line were analyzed with unpaired Student’s t-test. **(E)** Schematic of the experimental workflow for cytokine array analysis created in BioRender.com. **(F)** Cytokine array heatmap depicting fold changes in common SASP factors and immunomodulatory proteins after 7, 14, and 21 days of PMCi treatment in E98 senescent cells**. (G)** Gene Set Enrichment Analysis (GSEA) of RNA-seq data comparing non-responsive U251-MG cells treated with PMCi for 7 days versus untreated control cells, displaying normalized enrichment scores (NES) and corresponding p-values. **(E)** Heatmap of cytokine array data showing fold changes in cytokine levels between cells treated with PMCi alone and those treated with PMCi plus H151. **(H)** Colony formation assay comparing PMCi and H151 treatments (n = 2 independent experiments). Statistical analysis was performed using two-way ANOVA with Šidák's multiple comparisons test ($P < 0.05$). (**I**) Quantification of cell coverage in E98 cells following treatment with control medium (cMEM), conditioned medium from untreated cells after 7 days of incubation (C7CM), conditioned medium from senescent cells 7 days post-drug removal (S7CM), and PMCi. Data were analyzed using two-way ANOVA followed by Šidák's multiple comparisons test ($P < 0.05$). (**J**) Representative images of cell morphology (scale bar = 50 μM) and β-galactosidase assay (scale bar = 100 μM) in GBM cells treated with control conditioned medium (control c.m.) or PMC conditioned medium (PMC c.m.) for 7. Error bars represent the mean ± SEM.

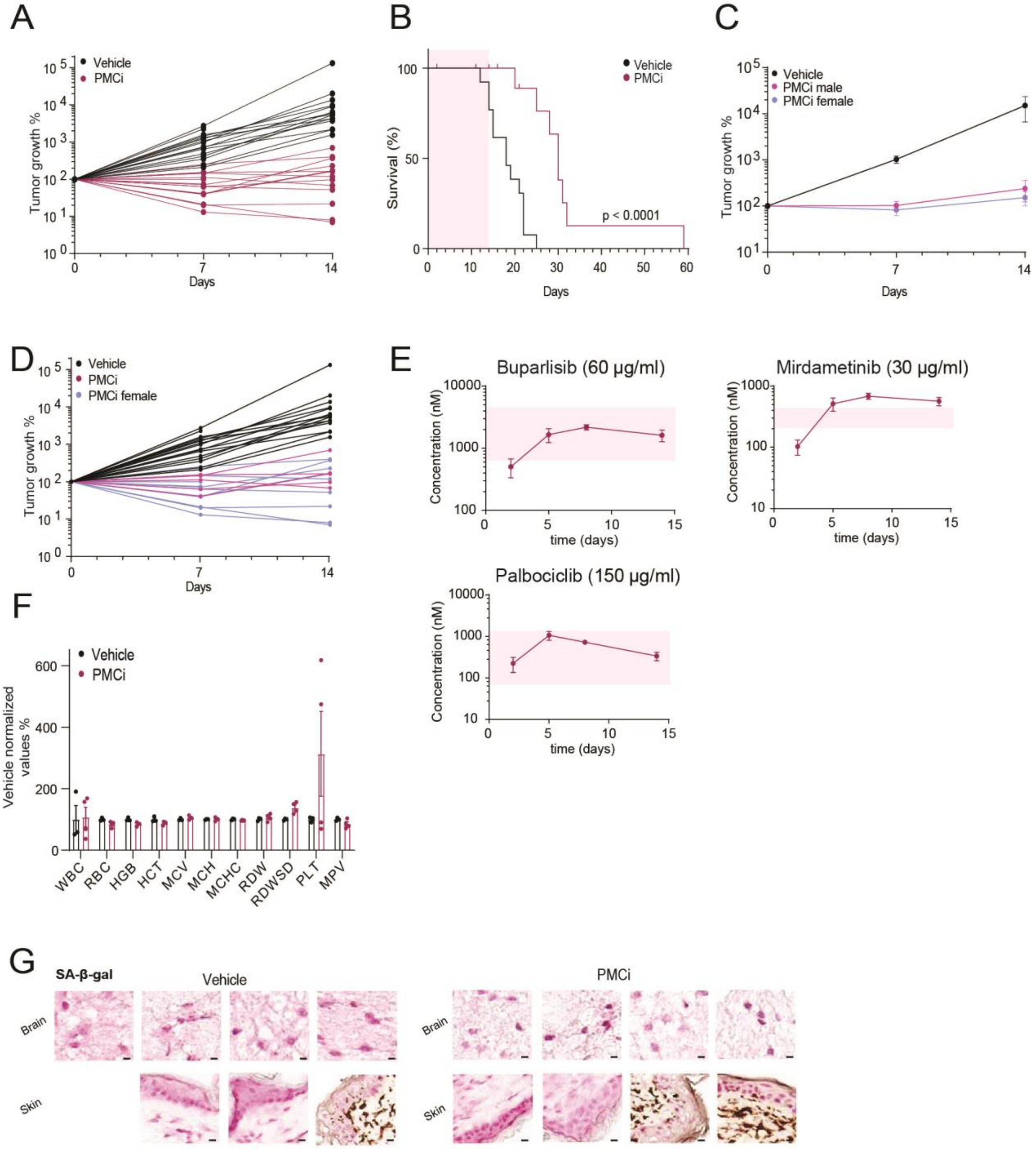


**Fig. S3. PMCi therapy is effective, tolerable and exhibits comparable effectiveness in both male and female mice.**

**(A)** tumor growth inhibition graph showing the individual values of the treated mice. **(B)** Kaplan-Meier survival plot. Mice with established orthotopic tumors were administered the PMCi treatment for 14 days. $\chi^2 = 13.11$, df = 1, $p < 0.01$, log-rank test. **(C)** Average tumor growth inhibition in male and female mice **(D)** and individual values of the same mice. **(E)** Drug plasma levels measured by LC-MS/MS in mice administered PMCi treatment via drinking water (drug concentration between brackets). The blue area represents the range between peak and trough plasma levels achieved in patients. **(F)** Normalized blood count values in mice treated for 14 days with PMCi (C). WBC : white blood cells, RBC: red blood cells,

HGB: hemoglobin, HCT: hematocrit, MCV: mean corpuscular volume, MCH: mean corpuscular hemoglobin, MCHC: mean corpuscular hemoglobin concentration, RDW: red cell distribution width, RDWSD: red cell distribution width standard deviation, PLT: platelet count, MPV: mean platelet volume. **(G)** Representative β-galactosidase-stained images of tissues from treated and untreated mice. Error bars: mean ± SEM.

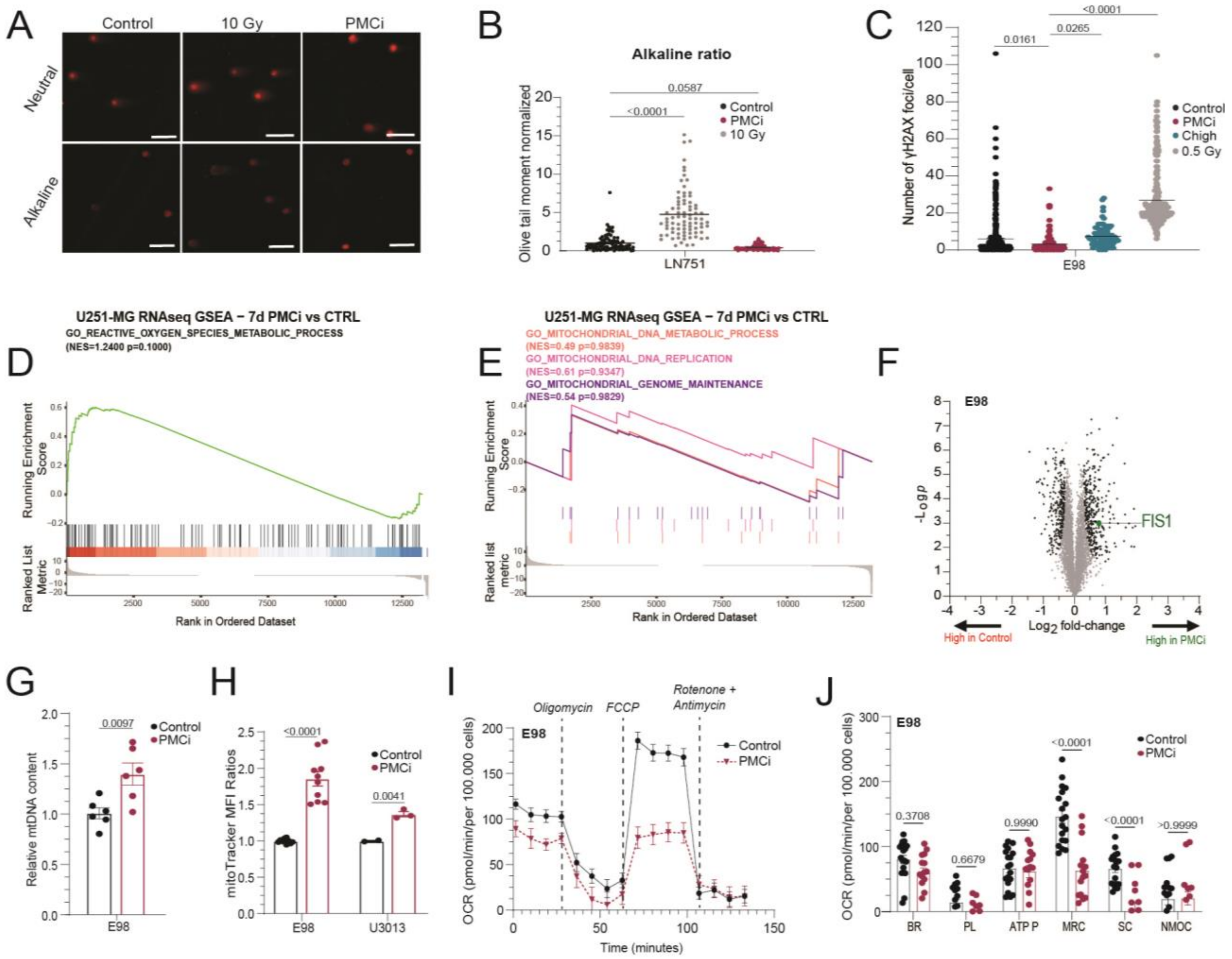


**Fig. S4. PMCi senescent cells show a decline in mitochondrial fitness despite the increased mass and gene expression.**

(A) Representative images from neutral and alkaline comet assays of untreated, irradiated, and PMCi-treated LN751 cells, visualized using fluorescence microscopy with a 20x objective and propidium iodide (PI) staining (scale bar = 20 μm). **(B)** Quantification of the olive tail moment in the alkaline comet assay, normalized to control values. Data from n = 2 independent experiments were analyzed by one-way ANOVA with Tukey's multiple comparison test. **(C)** Quantification of γH2AX foci in E98 GBM cells following treatment with PMCi or 1000 nM palbociclib (Chigh). Analysis was performed using one-way ANOVA with Tukey's multiple comparison test. **(D)** Gene Set Enrichment Analysis (GSEA) of RNA-seq data comparing U251-MG cells treated with PMCi for 7 days to untreated controls. **(E)** GSEA results for various mitochondrial-related gene sets, displaying normalized enrichment scores (NES) for each gene set, with distinct colors indicating different gene sets and corresponding p-values. **(F)** Volcano plot from proteomic analysis, highlighting changes in FIS1 expression following 7 days of PMCi treatment. Black dots indicate proteins significantly altered in senescent versus proliferative cells. **(G)** Quantification of mtDNA-to-nDNA copy number ratio after PMCi treatment, measured by qRT-PCR (n = 2 independent experiments). **(H)** Flow cytometric assessment of mitochondrial mass in E98 cells treated with PMCi, with median fluorescence intensity (MFI) ratios presented (n = 3 independent experiments). **(I)** Seahorse X-24 analysis comparing oxygen consumption rate (OCR) in control versus PMCi-induced senescent cells, including (J) quantification of basal respiration (BR), proton leak (PL), ATP production (ATP P), maximal respiratory capacity (MRC), spare respiratory capacity (SC), and non-mitochondrial oxygen consumption (NMOC).

Data from n = 3 independent experiments were analyzed with two-way ANOVA and Šídák's multiple comparisons test. Statistical analyses for panels **(G)** and **(H)** were conducted using unpaired Student's t-test. Error bars indicate mean ± SEM.

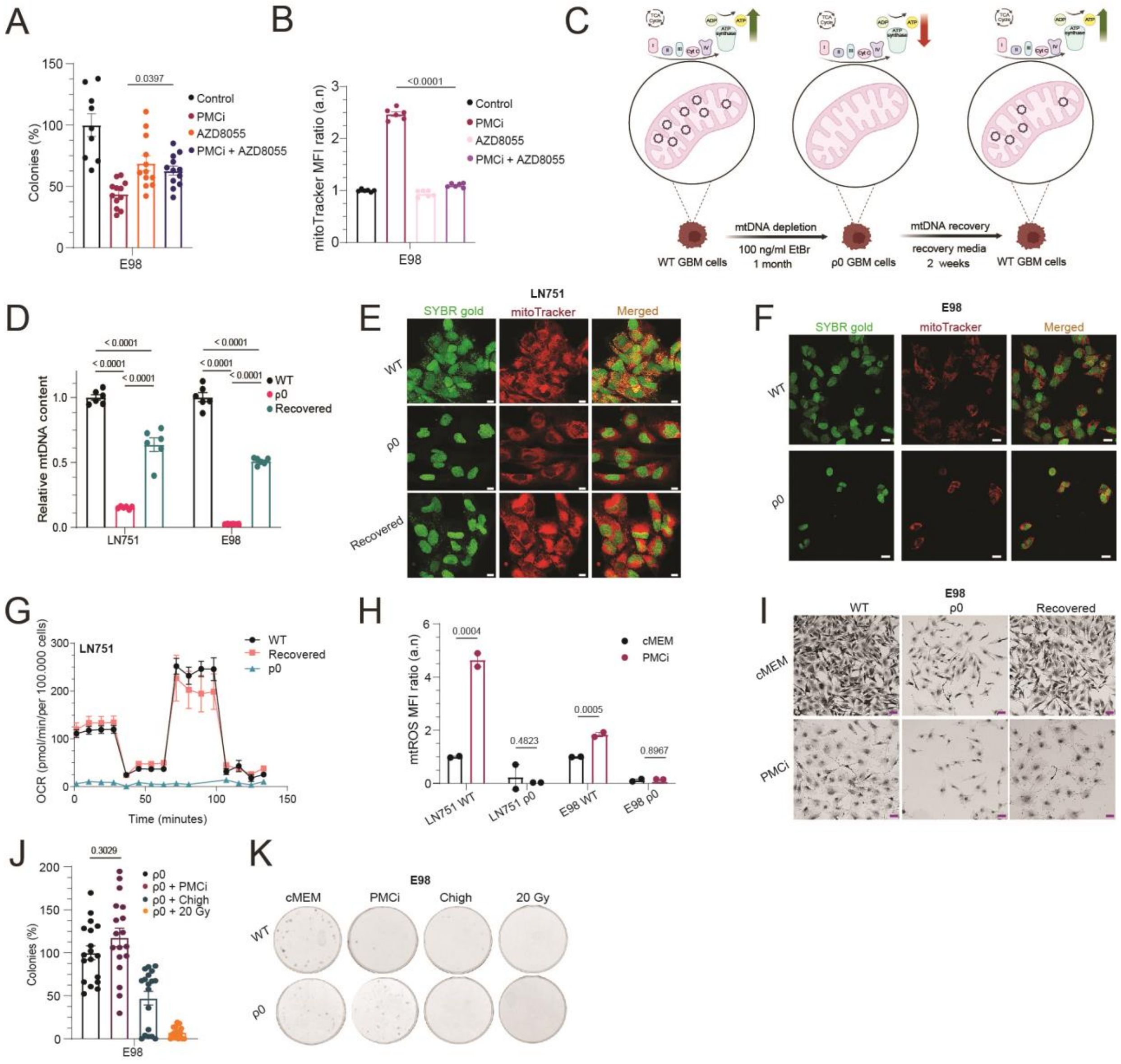


**Fig. S5. Depletion of functional mitochondria rescues the PMC-induced senescence characteristics.** **(A)** Colony formation assay of E98 cells treated with PMCi and 100nM of the mTOR inhibitor AZD8055. n=2 independent experiments. **(B)** MitoTracker-based flow cytometric detection of the same cells using MFI (median fluorescence intensity) ratios. **(C)** Schematic representation of the steps used for mtDNA depletion to generate ρ0 cells. Created in BioRender.com. **(D)** Quantification of the relative mtDNA-over-nDNA copy numbers using qRT-PCR. n=2 independent experiments. **(E)** Representative images showing mtDNA-depleted LN751 cells (ρ0) treated with low doses of ethidium bromide (EtBr) and mtDNA-recovered cells. Mitochondria were visualized using MitoTracker Deep Red, while mtDNA was stained with SYBR Gold. mtDNA appears as distinct punctate structures surrounding the nucleus. Scale bar: 15 µM. **(F)** Similar imaging was performed for E98 cells. Scale bar: 25 µM. **(G)** Seahorse X-24 metabolic analysis of oxygen consumption rate (OCR) in LN751 wild-type (WT), mtDNA-recovered, and ρ0 cells. **(H)** Flow cytometric detection of mitochondrial reactive oxygen species (mtROS) using MitoSOX, presented as median fluorescence intensity (MFI) ratios. **(I)** Morphological images of E98 control and ρ0 cells treated with a high dose of palbociclib (Chigh) and 20 Gy irradiation (scale bar = 50 µm). **(J)** Colony formation assay under the same treatment conditions, and **(K)** corresponding well plate images stained with crystal violet, comparing ρ0 cells with WT cells (n = 2 independent experiments). All bar graphs were analyzed using one-way ANOVA with Tukey's multiple comparison test. Error bars represent mean ± SEM.

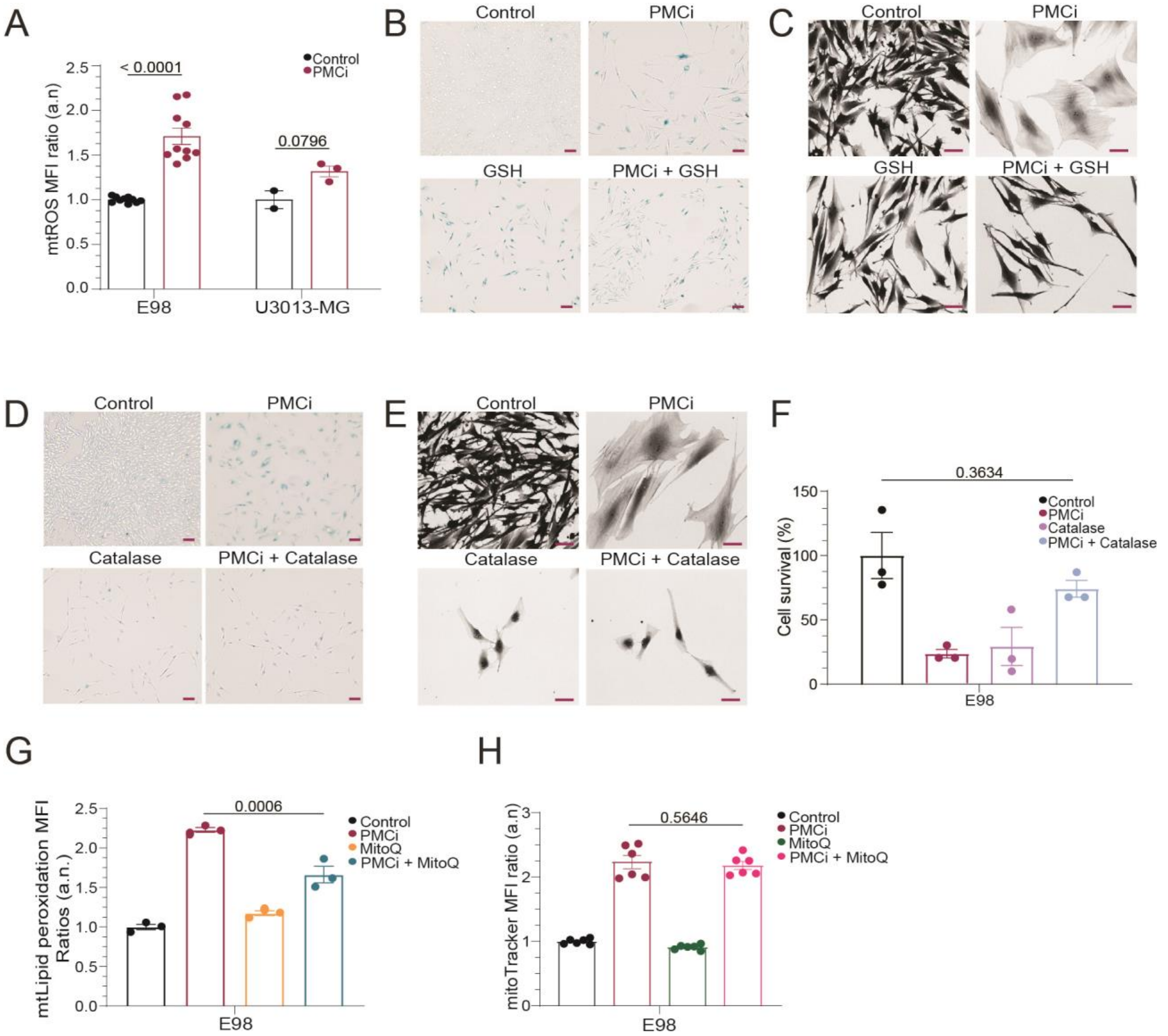


**Fig. S6. ROS scavengers rescue the PMCi- induced senescence phenotype and restore the proliferation arrest.**

**(A)** MitoSOX-based flow cytometric detection of mtROS, using MFI (median fluorescence intensity) ratios. Minimum n=2 independent experiments. Separate experiments per cell line were analyzed with unpaired Student's t-test. **(B)** Representative SA-β-gal photos (scale bar = 100μm) of LN751 cells treated with PMCi and 2mM glutathione (GSH) for 7 days. n=2 independent experiments. **(C)** Representative morphological photos (scale bar = 50Mμm) of LN-751 cells treated with PMCi for 7 days and GSH for the first 4 days of the PMCi exposure. **(D)** Representative SA-β-gal and **(E)** morphological images of LN751 cells treated with PMCi and 2000 U/mL catalase for 7 days. **(F)** colony formation assay using catalase. n = 1 performed triplicate. **(G)** MitoPerOx-based flow cytometric detection of lipid oxidation, using MFI ratios in E98 cells treated with 100 nM mitoQuinone (mitoQ) and PMCi and **(H)** MitoTracker-based flow cytometric detection, using MFI ratios, under the same conditions. n = 2 independent experiments. All bar graphs, except for **(A)**, were analyzed using one-way ANOVA with Tukey's multiple comparison test. Error bars represent mean ± SEM.

**Table S1. List of the antibodies used for western blot and simple western.**

| Antibody | Type | Species | Catalog No. | Supplier |
|---|---|---|---|---|
| p-AKT | Primary | Rabbit | 9271 | Cell signaling Technology |
| p-ERK | Primary | Rabbit | 4376 | Cell signaling Technology |
| p-Rb | Primary | Rabbit | 8180 | Cell signaling Technology |
| FOXM1 | Primary | Rabbit | 5436 | Cell signaling Technology |
| p-γH2AX | Primary | Rabbit | 9718 | Cell signaling Technology |
| a-rabbit HRP | Secondary | Goat | 7074 | Cell signaling Technology |
| a-rabbit 488 | Secondary | Goat | ab150077 | Abcam |

**Table S2. List of cytokines represented in the arrays shown in Figure 3E, Figure 8D, and Figure 9K.** Starting from the first columns the  cytokines are listed in the order they appear in the heatmaps of the main figures, from left to right.

| Cyokine list | | |
|---|---|---|
| Serpin E1 | VCAM-1 | BDNF |
| Leptin | TGF-α | BAFF |
| LIF | Thrombospondin-1 | Cystatin C |
| SHBG | RAGE | FGF-19 |
| Myeloperoxidase | RANTES | GDF-15 |
| Osteopontin | TNF-α | EGF |
| IL-19 | uPAR | CD14 |
| IL-22 | RBP-4 | Complement Component C5/C5a |
| IL-4 | Relaxin-2 | DPPIV |
| IL-5 | VEGF | G-CSF |
| GROα | Resistin | MIP-1α/MI |
| Growth Hormone | SDF-1α | MIP-3α |
| CXCL5 | MCP-3 | IL-34 |
| CD40 ligand | M-CSF | IP-10 |
| Adiponectin | IL-27 | IL-15 |
| Vitamin D B | IL-31 | IL-16 |
| ST2 | IL-10 | IL-1β |
| Lipocalin-2 | IL-11 | IL-1ra |
| MCP-1 | IFN-γ | CD30 |
| TARC | IGFBP-2 | EMMPRIN |
| PDGF-AA | Chitinase 3- | GM-CSF |
| PDGF-AB/B | Endoglin | MIP-3β |
| IL-23 | FGF-7 | MMP-9 |
| IL-24 | Cripto-1 | I-TAC |
| IL-6 | Angiopoietin-2 | |
| IL-8 | Angiopoietin-1 | |
| HGF | C-Reactive Protein | |
| ICAM-1 | FGF basic | |
| Fas Ligand | MIF | |
| Complement Factor | MIG | |
| Angiogenin | IL-32 | |
| Apolipoprotein A-I | IL-33 | |
| CD31 | IL-12 p70 | |
| TIM-3 | IL-13 | |
| TFF3 | IGFBP-3 | |
| TfR | IL-1α | |
| Pentraxin 3 | Flt-3 Ligand | |
| PF4 | Dkk-1 | |

**Table S3. List of the primers used for the amplification of mtDNA.**

| Gene | Primer sequence |
|---|---|
| GAPDH (nDNA) | Forward: TGGCCTCCAAGGAGTAAGACC<br>Reverse: CTGCCCCAGACCCTAGAATAAGA |
| short fragment (mtDNA) | Forward: ACTCTTTCACCCACAGCACC<br>Reverse: GGGGTTGAGGTCTTGGTGAG |
